# Longitudinal Exposomics in a Multiomic Wellness Cohort Reveals Distinctive and Dynamic Environmental Chemical Mixtures in Blood

**DOI:** 10.1101/2024.04.14.589329

**Authors:** Kalliroi Sdougkou, Stefano Papazian, Bénilde Bonnefille, Hongyu Xie, Fredrik Edfors, Linn Fagerberg, Mathias Uhlén, Göran Bergström, Leah JM Martin, Jonathan W. Martin

## Abstract

Chemical exposomes can now be comprehensively measured in human blood, but robust application of chemical exposomics in cohort studies requires knowledge of the longitudinal stability and interindividual variability of exogenous molecular profiles. Here we applied chemical exposomics to plasma of 46 adults, each sampled six times over two years in a multiomic wellness cohort. New chemicals were discovered, distinctive co-exposure patterns were observed, and intra-class correlation coefficients (ICC) for 519 confidently annotated substances are reported to support study design. Longitudinal stability of the chemical exposome (mean ICC 0.30) was significantly lower than the proteome, metabolome, lipidome or microbiome, and must be measured more frequently than other molecular profiles in health studies. Mixed-effects models nevertheless revealed significant associations between testosterone and perfluoroalkyl substances, and significant time-trends for low and high stability exposures alike. Complex exposome data structures were visualized and explored, demonstrating great potential for longitudinal exposomics in precision health research.

**Teaser:** The first cohort-level application of longitudinal exposomics revealed novel and dynamic co-exposures in blood of relevance to precision health.

## Introduction

The chemical exposome includes the cumulative sum of environmental chemical exposures throughout an individual’s life course. It includes exposure to natural and anthropogenic chemicals from external sources, such as inhalation of polluted air, intake of dietary substances and pharmaceuticals, and ingestion of contaminated food and water, but also includes internal exposure sources, such as the metabolic products of gut microbiota.(1,2) With recognition that our environment is dynamic over time, and that our susceptibility to disease changes over the life course, the exposome has always been imagined as a longitudinal endeavour that will require multiple measures of exposure.(1,2) For example, the external airborne chemical exposome has been shown to be highly dynamic for individuals over time, and variable among people, even for those living in the same geographical area.(3)

Comparatively little is known about the dynamics and variability of the chemical exposome in human blood. Sensitive methods for chemical exposomics have been described for blood plasma, including by multiclass targeted analysis(4,5) and combined targeted/untargeted analysis by liquid chromatography-high resolution mass spectrometry (LC-HRMS).(6,7) Measurement of the chemical exposome in blood is strategically important because the same sample can be further used for clinical testing, or for multiomic profiling of endogenous molecules whose levels may be impacted by the exposome, such as proteins or hormones.(8,9) Previous longitudinal studies of small molecules in blood have focused on the metabolome, with limited exploration of environmental chemicals in a few individuals for up to a few months.(9,10) Studies of the blood chemical exposome have yet to be reported in a longitudinal cohort, and consequently there are few measures of inter- and intra-individual variability for priority environmental substances. Such data could inform the number of sampling events to include in future exposome studies (i.e. to ensure statistical power), and longitudinal data are inherently necessary to differentiate short-term and chronic exposures.

Here we applied chemical exposomics to recurrent plasma samples of 46 healthy Swedish adults, each sampled six times over 2 years in a multiomic wellness profiling study. Using LC-HRMS for quantitative analysis of 83 multiclass targeted analytes (78 priority environmental contaminants and 5 steroid hormones) and parallel untargeted discovery of environmental chemicals and endogenous metabolites,(7) here we report the inter- and intra-individual variability for hundreds of molecular environmental exposures, including chemicals not previously detected in human blood. Intra-class correlation coefficients (ICCs) revealed that the stability of the chemical exposome was generally low, compared to parallel measures of plasma proteome, metabolome, lipidome, microbiome in the same participants. Repeated sampling of the same individuals over time permitted new exposure-types to be defined, and for rare and common co-exposures to be revealed of relevance to precision health. Mixed-effect modelling also revealed statistically significant temporal trends and exposome-metabolome interactions indicative of endocrine disruption.

## Results and Discussion

### Multiclass Targeted Analysis

Among all plasma samples of males and females, 57 out of the 83 target analytes were detected and quantified. These substances belonged to a diverse range of 14 chemical classes (**Table S4**, **Figure 1A-1B**), including fungicides, herbicides, insecticides (organophosphate and neonicotinoid), flame retardants, PFAS, personal care products, pharmaceuticals, plasticizers, phytoestrogens, dietary additives, polycyclic aromatic compounds, a nicotine metabolite, and endogenous steroid hormones. For further statistical and multivariate analysis, only target analytes with DF > 10% were considered, including 34 analytes from 9 chemical classes (**Figure S1**). There were few differences by sex, but comparing mean plasma concentrations of males (n=138) and females (n=138), significant differences were identified for parabens, as well as certain PFAS and hormones (**Figure 1A-1B**, Student’s t-test, two-tailed, Bonferroni corrected p-values). Methylparaben and propylparaben, which are used in cosmetics, had significantly higher levels in females (methylparaben: males 0.60 ng/mL, females 1.5 ng/mL, p < 0.001; propylparaben: males 0.06 ng/mL, females 0.31 ng/mL, p < 0.01, **Figure 1A**), consistent with a previous study in urine(11). Conversely, females had significantly lower concentrations of several PFAS (**Figure 1B**), consistent with known routes of PFAS elimination during pregnancy, breastfeeding, and menstruation,(12–15) including for linear PFHxS (males 1.4 ng/mL, females 0.90 ng/mL, p < 0.001), linear PFHpS (males 0.25 ng/mL, females 0.16 ng/mL, p < 0.001), branched PFHpS (males 0.033 ng/mL, females 0.017 ng/mL, p < 0.001) and branched PFOS (males 3.11 ng/mL, females 2.40 ng/mL, p < 0.001). As expected, mean testosterone was significantly higher in males (3.2 ng/mL) than females (0.18 ng/mL, p < 0.001) (**Figure 1A**), with levels in reference ranges for healthy adults in comparable age groups.(16)

**Figure 1.**
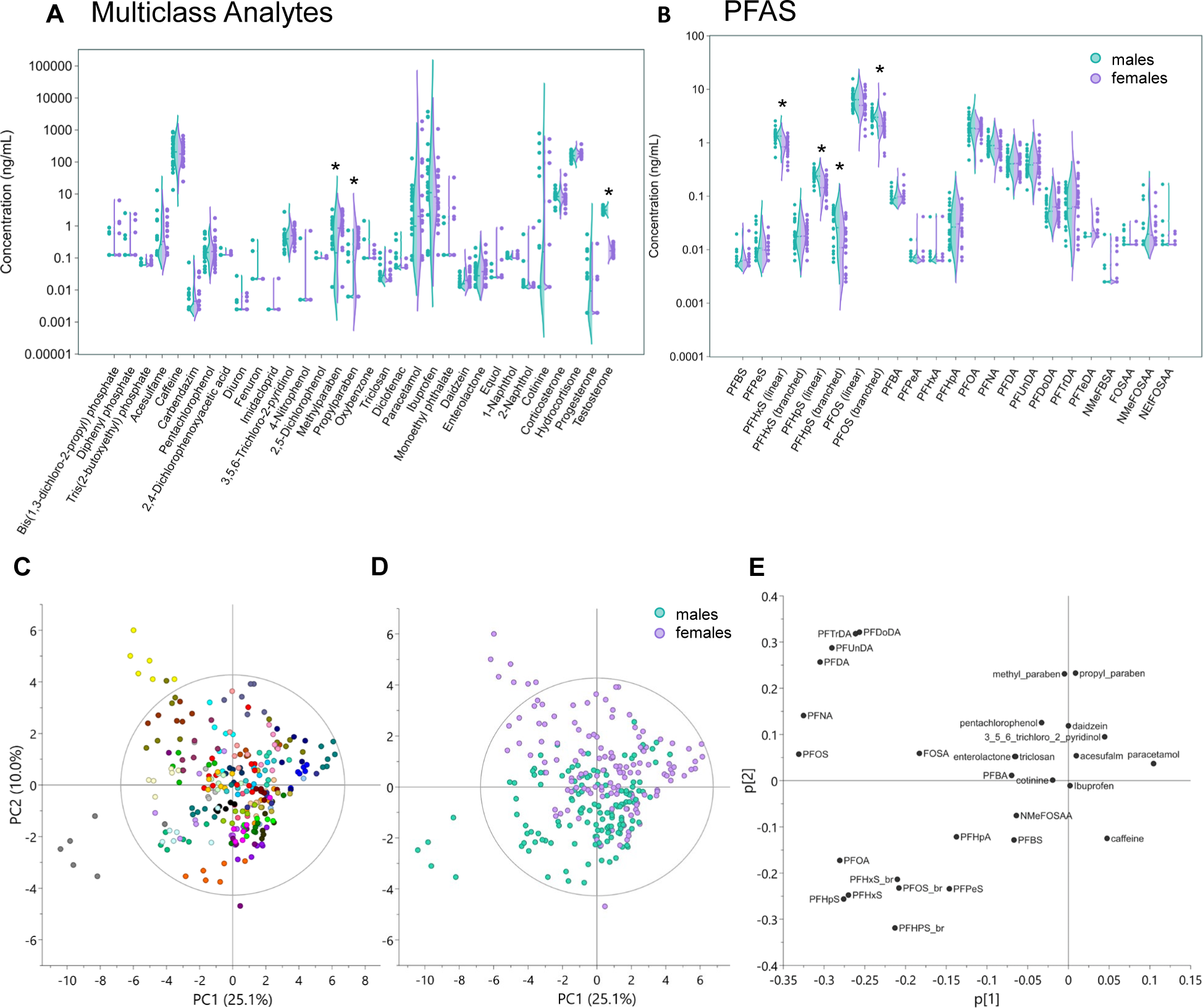
Targeted chemical exposome and individual variation in S3WP participants. Panels (A) and (B) show violin plots of average analyte concentrations per individual (log_10_ scale) for targeted multiclass analytes and targeted perfluoroalkyl and polyfluoroalkyl substances (PFAS), respectively, with color coding according to males (green, n = 23) and females (purple, n = 23). Dashed lines on violin plots indicate mean concentration for each analyte in each sex. Asterisks on top of violin plots indicate the analytes with a significant different concentration between males and females (Bonferroni corrected p < 0.01; Student’s t-test, two-tailed, n= 138 for each sex). In panel (C) and (D) the 6 sampling visits of each individual are shown in principal component analysis (PCA) scores plots (3 components; R2Xcum = 43.4%, Q2cum = 29.8%) with samples color coded by participant ID (C) and sex (D); PCA loadings of the targeted analytes are shown in panel (E). In the PCA plots, only analytes with a detection frequency > 10% are shown, and excluding steroid hormones.

After excluding the detected sex hormones (hydrocortisone, corticosterone, progesterone and testosterone), PCA was applied to broadly visualize and detect sources of intra- and inter-individual variability of environmental exposures (30 detected targeted analytes) over the 2-year period in this adult population (**Figure 1C-1E**). In the PCA scores plot colored by S3WP participant ID (**Figure 1C**), the 6 samples from each individual were often grouped together, indicating relative stability of the target exposome over 2 years. Notably, two individuals were consistently outliers relative to the study population, showing particularly high levels of certain PFAS (i.e. C10-C13 perfluoroalkyl carboxylates for female W0043 colored in yellow; C5-C8 perfluoroalkyl sulfonates for male W0074, colored in gray, **Figure 1E**). Partial separation between males and females along PC2 (10% of variation, **Figure 1D**) was mostly explained by sex-specific exposures to parabens and PFAS (**Figure 1E**), consistent with statistical sex differences noted above.

### Untargeted Molecular Discovery

Among all 276 plasma samples, a total of 129,547 unique untargeted features were detected across ESI+ and ESI-analyses after blank filtering and removal of redundant features; thus, targeted analytes represented only 0.04% of the overall molecular dataset. Substantially fewer features were detectable in any individual person (mean 73,426 features per individual, range 67,405 – 87,159 features, **Figure S2**). Moreover, only 20,520 features (16% of total dataset) were detected in all individuals in at least one visit, somewhat lower than for the targeted analytes (i.e. 26% of targeted analytes were detected in all individuals for at least one time point). These results indicate that a high proportion of ‘molecular dark matter’(17) in plasma is unique to individuals, possibly representing unique environmental exposures, and/or unique biological response at the metabolome level. This finding is consistent with a recent observation that gut microbiota composition (i.e. internal exposome) explains the majority of variance (i.e. 58%) in individual human plasma metabolites,(18) although other sources of environmental exposure have yet to be similarly examined. In untargeted studies of the exposome or the metabolome, it is therefore predictable that the complexity of molecular datasets will increase with larger sample sizes.

By automated matching to open access libraries (based on MS1 accurate mass and MS2 spectra) and after manual curation of all untargeted dataset annotations, a total of 462 high confidence structural candidates were assigned (i.e. with at least Level 2a identification confidence(19)) in ESI+ and ESI-, corresponding to a 0.4% annotation rate; this total does not include the targeted analytes, which are considered confirmed at Level 1 confidence. Combining all the Level 1 identified chemicals (targeted analytes and untargeted molecular discoveries confirmed by reference standard(19); see next section), and all Level 2 high confidence annotations, resulted in a total of 519 substances, including 343 environmental chemicals, 162 endogenous metabolites and 14 substances with ambiguous classification (**Table S5**). The 343 environmental chemicals were divided into 11 subclasses, including foods and additives, drugs and their metabolites, industrial chemicals, PFAS, flame retardants, plasticizers, personal care products, plant metabolites and natural products (**Figure S3**), and is hereafter referred to as the “chemical exposome” to distinguish it from the endogenous metabolites and/or substances with ambiguous classification. The 162 endogenous metabolites were divided into 7 subclasses, including fatty acids and derivatives, bile acids, amino acids and hormones. We acknowledge that a strict separation between chemical exposome and endogenous substances leaves certain ambiguities, including for the reason that the human metabolome includes the endogenous metabolomes of other species consumed in our diet.(20)

### Confirmation of Selected Structural Annotations

To confirm the Level 2 molecular annotations from untargeted analysis, authentic standards (>98% purity) were obtained for a set of 25 analytes, including 3 endogenous and 22 environmental substances. Confirmations were successful for 20 analytes (80% success rate, **Tables 1 and S6, Figures S4-S21**), indicating an effective untargeted acquisition and data-processing workflow for molecular discovery. Several unexpected and widespread environmental exposures were thereby confirmed (Level 1), including rubber additives, plasticizers, industrial chemicals and personal care products **(Table 1)**. When a confirmed analyte was also detectable in the pooled Swedish reference plasma, and its concentration could be quantified with a standard addition curve (0-10 ng/mL range), reference standardization(21) was used to semi-quantify these discovered chemicals in the individual samples (**Table 1**). Selected examples of confirmed molecular discoveries are discussed below.

**Table 1.**
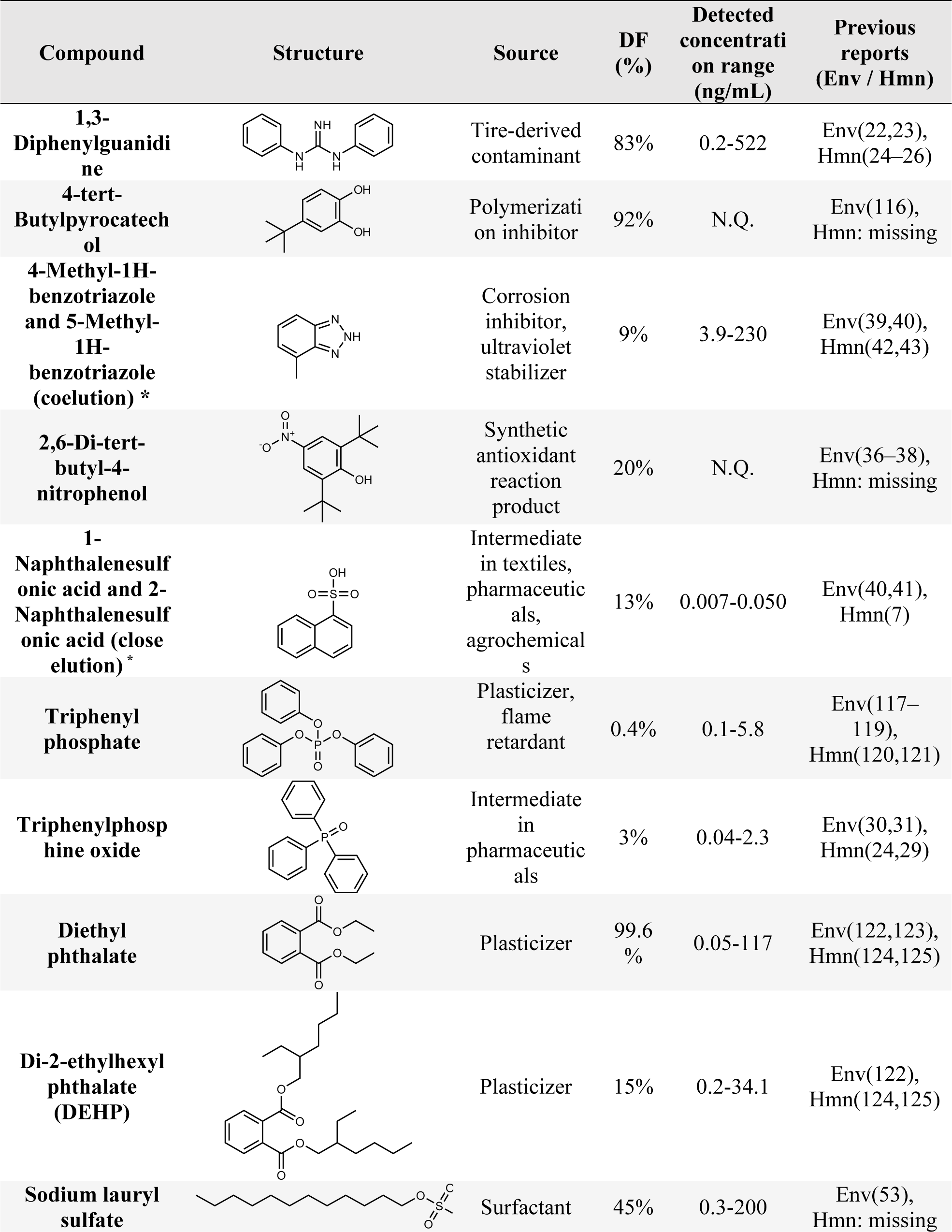
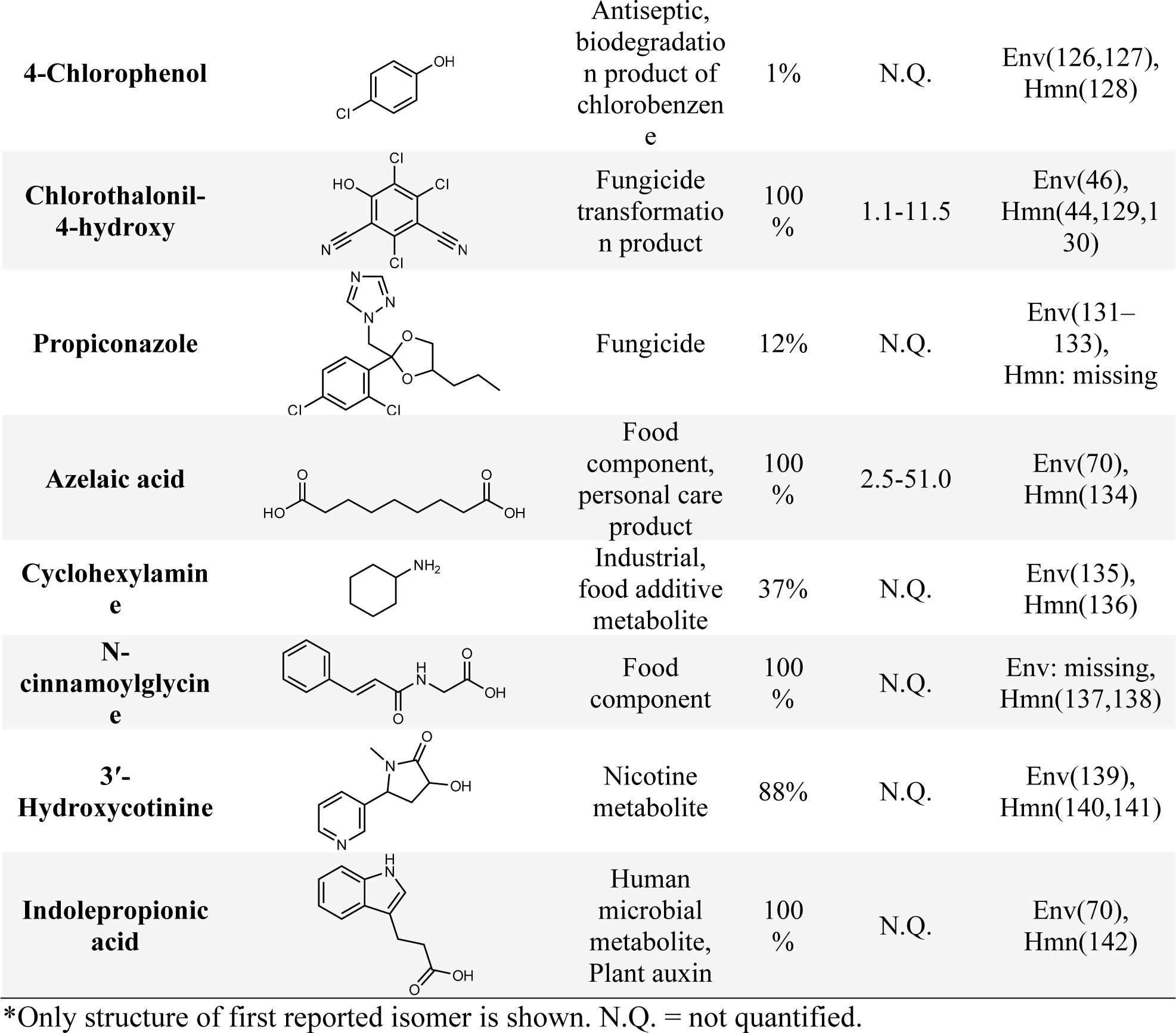
Untargeted molecular discoveries in plasma samples confirmed at confidence Level 1. The table shows chemical name, structure, sources, detection frequency (DF), concentration range among detections (semi-quantification), and previous reports in environmental (Env) or human (Hmn) samples searched in CAS SciFinderⁿ(115).

| Compound | Structure | Source | DF (%) | Detected concentration range (ng/mL) | Previous reports (Env / Hmn) |
| --- | --- | --- | --- | --- | --- |
| 1,3-Diphenylguanidine |  | Tire-derived contaminant | 83% | 0.2-522 | Env(22,23), Hmn(24-26) |
| 4-tert-Butylpyrocatechol |  | Polymerization inhibitor | 92% | N.Q. | Env(116), Hmn: missing |
| 4-Methyl-1H-benzotriazole and 5-Methyl-1H-benzotriazole (coelution) * |  | Corrosion inhibitor, ultraviolet stabilizer | 9% | 3.9-230 | Env(39,40), Hmn(42,43) |
| 2,6-Di-tert-butyl-4-nitrophenol |  | Synthetic antioxidant reaction product | 20% | N.Q. | Env(36-38), Hmn: missing |
| 1-Naphthalenesulfonic acid and 2-Naphthalenesulfonic acid (close elution) * |  | Intermediate in textiles, pharmaceuticals, agrochemicals | 13% | 0.007-0.050 | Env(40,41), Hmn(7) |
| Triphenyl phosphate |  | Plasticizer, flame retardant | 0.4% | 0.1-5.8 | Env(117-119), Hmn(120,121) |
| Triphenylphosphine oxide |  | Intermediate in pharmaceuticals | 3% | 0.04-2.3 | Env(30,31), Hmn(24,29) |
| Diethyl phthalate |  | Plasticizer | 99.6% | 0.05-117 | Env(122,123), Hmn(124,125) |
| Di-2-ethylhexyl phthalate (DEHP) |  | Plasticizer | 15% | 0.2-34.1 | Env(122), Hmn(124,125) |
| Sodium lauryl sulfate |  | Surfactant | 45% | 0.3-200 | Env(53), Hmn: missing |
| **4-Chlorophenol** |  | Antiseptic,<br>biodegradation<br>product of<br>chlorobenzene | 1% | N.Q. | Env(126,127),<br>Hmn(128) |
| **Chlorothalonil-4-hydroxy** |  | Fungicide<br>transformation<br>product | 100<br>% | 1.1-11.5 | Env(46),<br>Hmn(44,129,130) |
| **Propiconazole** |  | Fungicide | 12% | N.Q. | Env(131-133),<br>Hmn: missing |
| **Azelaic acid** |  | Food<br>component,<br>personal care<br>product | 100<br>% | 2.5-51.0 | Env(70),<br>Hmn(134) |
| **Cyclohexylamine** |  | Industrial,<br>food additive<br>metabolite | 37% | N.Q. | Env(135),<br>Hmn(136) |
| **N-cinnamoylglycine** |  | Food<br>component | 100<br>% | N.Q. | Env: missing,<br>Hmn(137,138) |
| **3'-Hydroxycotinine** |  | Nicotine<br>metabolite | 88% | N.Q. | Env(139),<br>Hmn(140,141) |
| **Indolepropionic acid** |  | Human<br>microbial<br>metabolite,<br>Plant auxin | 100<br>% | N.Q. | Env(70),<br>Hmn(142) |

\*Only structure of first reported isomer is shown. N.Q. = not quantified.

#### Rubber additives

The tire-derived substance, 1,3-diphenyl guanidine (**Table 1, Figure S4**) was recently reported to be the most frequently detected tire-derived contaminant in air(22) and indoor dust(23) from around the globe and was confirmed here in 83% of samples. Reports of 1,3-diphenyl guanidine in human biofluids remain rare.(24–26) A related substance was confirmed, 4-tert-butylcatechol (TBC, **Table 1, Figure S7**), detectable in 92% of samples despite no previous reports in human biofluids. This substance is reported as a contact allergen,(27) and is used as an additive polymerization inhibitor in commercial monomers, such as styrene, and used in the rubber, paint and petroleum industry.(27,28).

#### Industrial chemicals

Triphenyl phosphine oxide (TPPO) is a widely used synthetic intermediate in pharmaceutical products(29) and was detected here in 3% of samples (**Table 1, Figure S5**). TPPO has begun to be widely detected in indoor air, dust and aquatic systems,(30,31) but has only rarely been detected in human samples.(24,29) Other confirmed industrial intermediates were the related isomers 1- and 2-naphthalene sulfonate (**Table 1, Figure S12**), used in textile, pharmaceutical and agrochemical production(32,33). These two substances are known to have low biodegradability(32,33) and neither has been previously confirmed in human biofluids (2-naphthalene sulfonate has been reported previously at Level 2(7)) but were detected in 13% of samples here.

We also discovered and confirmed 2,6-di-tert-butyl-4-nitrophenol (DBNP, **Table 1, Figure S6**), a known transformation product of 2,6-di-tert-butyl-4-methylphenol (BHT).(34) BHT is a widely used synthetic antioxidant added to polymers, foods and cosmetics, that has been detected in the environment and in human biofluids and tissues in several countries,(35) but DBNP has not previously been reported in human biofluids, although it has been detected in the environment(36,37) and in plastic food packaging as a “non-intentionally added substance”.(38) Its detection in 20% of Swedish samples here deserves further attention due to its biological persistence and toxic potential through uncoupling of oxidative phosphorylation.(37) Two benzotriazole isomers, 4-methyl- and 5-methyl-1H-benzotriazole, were also confirmed and semi-quantified in 9% of samples (**Table 1, Figure S10**). These benzotriazoles are high-production volume chemicals mainly applied as corrosion inhibitors and ultraviolet stabilizers, widely detected in environmental matrices(39–41) but also in a few studies in human samples.(42,43)

#### Pesticides

Chlorothalonil-4-hydroxy was confirmed and semi-quantified in 100% of plasma samples (1.1-11.5 ng/mL, median 3.7 ng/mL; **Table 1, Figure S9**). This finding is consistent with a targeted study of pregnant Swedish women (1997–2015) where it was also detected in all samples (median 4.1 ng/mL),(44) yet it remains rarely monitored in humans. It is a known transformation product of the fungicide chlorothalonil, which has not been permitted for agricultural use in Sweden since the 1990s,(45) and has been banned in the EU since 2019 due to its carcinogenic properties and risk to fish and amphibians.(46) Chlorothalonil-4-hydroxy is considered more toxic,(47) more persistent and more mobile in soil(48) than the parent pesticide, thus its widespread presence in blood deserves attention in exposome studies.

#### Personal care products

Sodium lauryl sulfate was confirmed and detected in 45% of samples (**Table 1**, **Figure S8**), and has not previously been reported in human biofluids to our knowledge despite being a major surfactant in shampoos.(49,50) It is known to be absorbed into the blood stream in animal models,(51) and may also be inhalable during shampoo use.(52) This surfactant is rarely monitored and remains unregulated due to its biodegradable nature, but its risks have been debated.(53)

### Longitudinal Stability of the Chemical Exposome

The final combined dataset of 519 annotated substances (Level 1 and 2) was used to examine the intra- and inter-individual variation of exposure in this 2-year study with six sampling points. For each substance we calculated the ICC (**Table S7**), a non-dimensional ratio of the inter-individual variance to the total variance (i.e., the sum of the inter- and intra-individual variance).(54) In context of exposome research, ICCs are instructive for study-design as they describe the extent to which individuals retain their rank-order in a study population(55) with repeated measurements of exposure over time. Higher ICCs correspond to more stable exposures, that can therefore be measured fewer times throughout the life course, whereas lower ICCs correspond to less stable exposures that may need several repeated measurements to accurately classify exposure over the life course. ICCs range from 0 to 1, with values < 0.40 corresponding to poor reproducibility of repeated measurements, 0.40 to 0.75 is considered fair to good, and >0.75 indicates excellent reproducibility.(54)

The majority of annotated substances in plasma (306 of 519 analytes) had ICCs < 0.40 (**Figure 2A**), and the mean ICC was significantly higher for endogenous metabolites (0.40) than for the chemical exposome (0.30, Student’s t-test, two-tailed, p < 0.001) (**Figure 2B**). Moreover, while the ICCs for endogenous metabolites were normally distributed, the chemical exposome showed denser distributions towards lower and higher ICC values. More specifically, 66% of ICCs were < 0.4 for the chemical exposome, compared to only 45% of ICCs for endogenous metabolites, and 10% of the ICCs were > 0.75 for the chemical exposome, compared to only 6% of ICCs for endogenous metabolites (**Figure 2B**). While the plasma chemical exposome and metabolome have highly stable components, these results mean that the majority of small molecules in plasma must be measured more than once over time to adequately represent an individual’s exposure.

**Figure 2.**
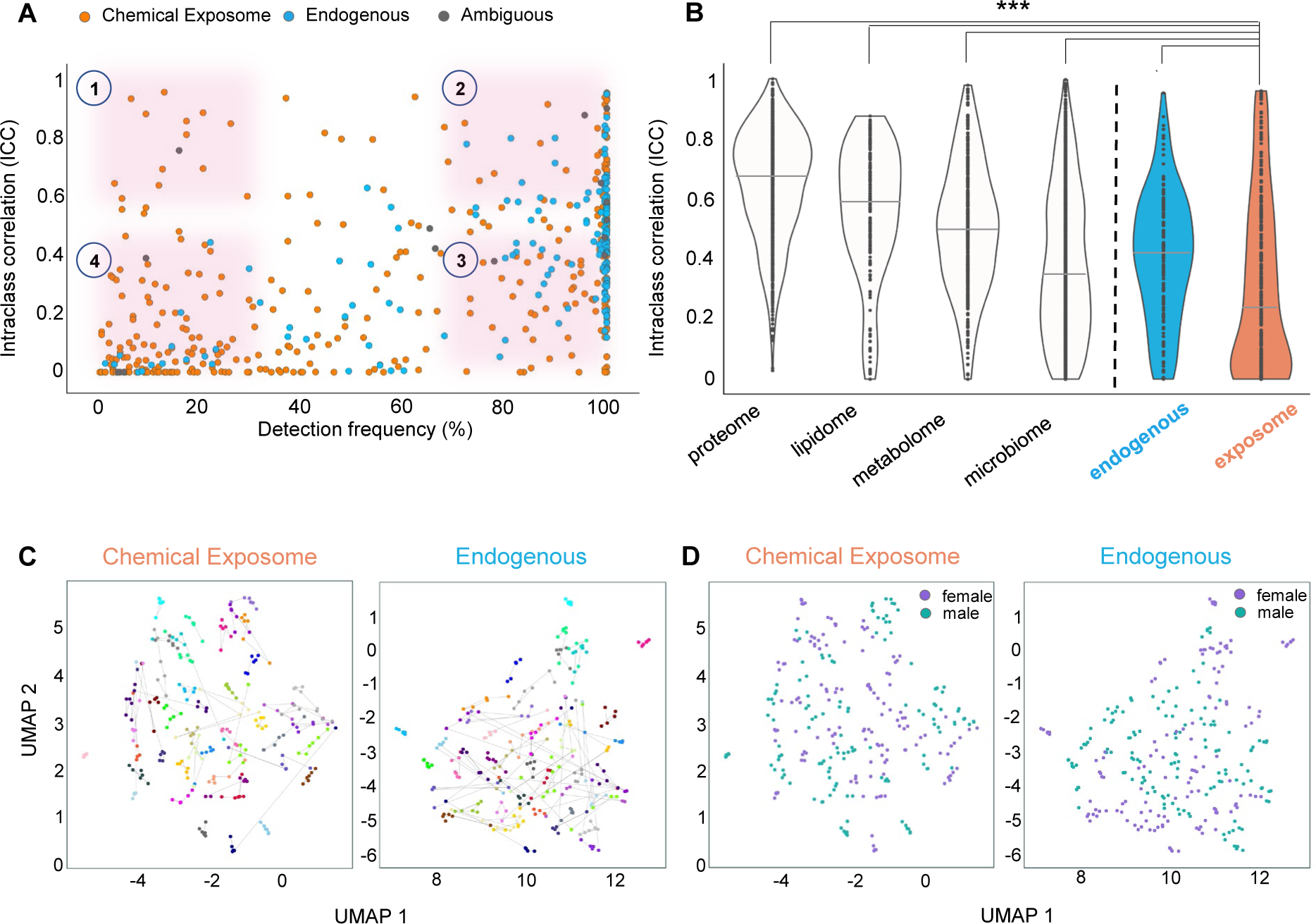
Longitudinal stability of the chemical exposome and comparison to other molecular profiles. Panel (A) shows the calculated intraclass correlation coefficients (ICC) of the 519 annotated substances (Level 1 and Level 2) along with their detection frequency (DF), for the chemical exposome (orange, n = 343), endogenous metabolites (blue, n =162) and compounds with ambiguous origin (gray, n = 14). Four regions of interest are highlighted on the plot: up left (Type 1; rare-stable, low DF, high ICC), up right (Type 2; common-stable, high DF, high ICC), bottom right (Type 3; common-unstable, high DF, low ICC), bottom left (Type 4; rare-unstable, low DF, low ICC). In panel (B) the ICC data are presented in violin plots separately for the chemical exposome and the endogenous metabolites as well as for the proteome, lipidome, metabolome and microbiome from Tebani et al.(57), including only individuals present in the exposome study. In panel (B) solid lines show median ICC values and asterisks indicate significantly different ICC means between the chemical exposome and endogenous metabolites (p < 0.001; Student’s t-test, two-tailed) and between the chemical exposome and all other molecular profiles (p < 0.001; one-way ANOVA and Tukey’s test). Panels (C) and (D) show global profiles across visits applying uniform manifold approximation and projection (UMAP) for the chemical exposome and the endogenous metabolites color coded by participant ID with gray lines connecting samples representing different visits of the same individual (C) or color coded by sex (D).

To conclusively link environmental exposures to disease etiology, an objective in exposomics is to identify key perturbations in molecular networks at the metabolome, proteome and gene expression levels.(56) In longitudinal exposome studies, this might be addressed through multiomics, but the relative dynamics of each molecular dataset must be considered at the study design stages. Thus, here we compare our results to those of Tebani et al. who reported ICCs for several multiomic profiles in the same study participants.(57) We recalculated the ICCs of Tebani et al. to only include individuals with paired chemical exposome measurements (see Supporting Information), and compared to the relatively low stability of the chemical exposome (mean ICC = 0.30), significantly higher mean ICCs were evident for the plasma proteome (0.65), lipidome (0.55), metabolome (0.50), and gut microbiota (0.38) (**Figure 2B**; one-way ANOVA, Tukey’s test p < 0.001). These results emphasize the importance of repeated measures of the chemical exposome in epidemiological studies to minimize exposure misclassification, and in longitudinal multiomic studies the exposome should be measured as frequently, or more frequently, than other biomolecular profiles.

The gut microbiome had the most similar distribution of ICCs in comparison to the chemical exposome, with overall lower reproducibility, and similar high density at lowest values (**Figure 2B**). Like the chemical exposome, gut microbiota is shaped by life course exposures, including diet, disease history and medication, as well as by intrinsic factors such as age and host genetic variation.(18) As discussed above, the known link between gut microbiota and plasma small molecules(18) could partially explain similar ICC distributions for gut microbiome and the plasma chemical exposome.

Despite relatively low temporal stability overall, the chemical exposome has stable components. This was highlighted by UMAP analysis, allowing the relative variation of individual exposomes over time to be visualized relative to the study population (**Figures 2C and 2D**). The visits of each individual, colored by participant ID and connected by lines, demonstrate that the complex chemical exposome profile of most individuals has stable factors over time (**Figures 2C**). However, individual variability was evident both for the chemical exposome and endogenous metabolites, with some individuals having remarkably unique and stable profiles, and others displaying higher variability between visits (**Figure 2C**). UMAP visualization of metabolome stability in Tebani et al.(57) showed similar results. Notably, UMAP projections of the chemical exposome and endogenous metabolites did not indicate any sex differences (**Figure 2D**).

### Longitudinal Trends

Temporal trend studies of human exposure to environmental pollutants are normally conducted with cross-sectional sampling designs covering decades (i.e. different people analyzed in various years). (58,59) Thus, the high-temporal resolution of the current longitudinal study of 46 individuals presented a unique opportunity to examine if temporal trends could be detected in a comparatively short period of only two-years. Using linear mixed-effects regression models, which account for the repeated sampling of individuals, we detected statistically significant declining temporal trends for this adult population. Focusing on quantified targeted analytes with high detection frequencies, here we report two such examples: a relatively stable analyte with high ICC (pentachlorophenol, ICC 0.83, 98% DF) and another with particularly low stability (3,5,6-trichloro-pyridinol, ICC 0.25, 97% DF**) (Figure 3**). For pentachlorophenol, concentrations decreased over time with a rate constant of −0.19 years^-1^ (95% CI: −0.078, −0.29), corresponding to a disappearance half-life of 3.7 yrs (95% CI: 2.4-8.8 yrs). This value compares remarkably well to a reported population disappearance half-life of 3.6 yrs calculated from a global meta-analysis of blood pentachlorophenol concentrations compiled over 30 years (1978–2008)(60). Sweden was the first country to control the use of this pesticide in 1978, and many countries have since followed suit(60), thus continued declining concentrations observed here in Swedish adults is positive news for this carcinogenic and endocrine active substance.(60) For 3,5,6-trichloro-pyridinol, concentrations decreased over time with a rate constant of −0.26 years^-1^ (95% CI: −0.055, −0.46), corresponding to a disappearance half-life of 2.7 yrs (95% CI: 1.5-12.6 yrs). The wider confidence interval is reflective of the lower stability for this analyte, relative to pentachlorophenol, but the statistically significant result is particularly noteworthy as it demonstrates the value of repeated exposome measurement in longitudinal study designs such as this. The result is moreover of interest because 3,5,6-trichloro-pyridinol is a metabolite of chlorpyrifos, a neurotoxic insecticide that was effectively banned or phased-out in the EU and United States by 2020(61). Hites recently reviewed the evidence from human biomonitoring of 3,5,6-trichloro-pyridinol in the United States and Sweden, and although mean concentrations were declining it was concluded that the statistical support for the trends were weak(61). These examples demonstrate that longitudinal exposome studies will be a rich source of data for chemical regulators, as well as health scientists.

**Figure 3.**
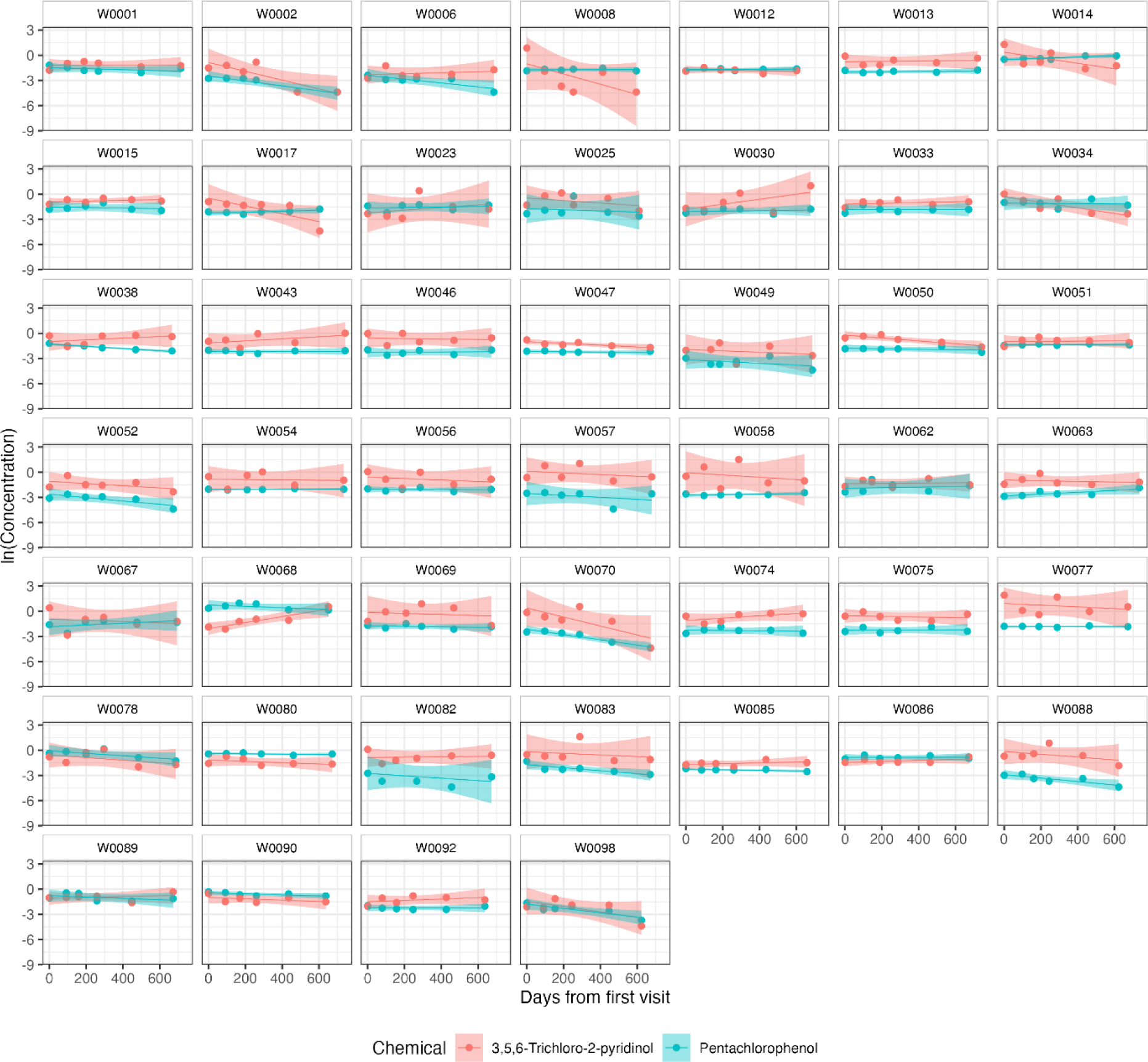
Example temporal trends of two targeted analyte concentrations in plasma of participants. Pentachlorophenol (blue) and 3,5,6-trichloro-2-pyridinol (red) concentrations (natural log) are shown in each individual’s six samples over the two-year study. Simple linear regression lines with 95% confidence intervals are shown, but statistical significance of the population temporal trend was evaluated in mixed-effects models.

### Longitudinal Exposure Types

A plot of each chemical’s ICC versus sample detection frequency (**Figure 2A**) allowed for new insights to chemical exposure characterization that would not have been evident without repeated sampling in this longitudinal study. Four exposure-Types (inset #1-4, **Figure 2A)** are defined and discussed in this context. First, we categorize Type 1 exposures as rare-stable (i.e. detection frequency < 20%, ICC 0.6-1). Although such exposures are not detectable in most people, and could therefore be easily overlooked in cross-sectional study designs, these exposures may be of high relevance to the small fraction of individuals with consistently elevated exposure over the life course. These exposures are therefore of high relevance in a precision health context, as they likely arise from unique behaviors, lifestyle factors or occupational exposures that could be mitigated if identified. An example was the industrial intermediate triphenyl phosphine oxide (ICC 0.64, 3.2% DF; Level 1) which was only detected in 9 samples overall, but consistently in all 6 samples of individual W0015. Other examples included NEtFOSAA (ICC 0.93, 7% DF; Level 1) (consistent for individual W0008), a PFOS precursor with exposure sources linked to food packaging(62) and the drug citalopram (ICC 0.88, 9% DF; Level 2) (consistent for individual W0090), which acts as a selective serotonin reuptake inhibitor(63).

Type 2 exposures were categorized as common-stable (i.e. detection frequency > 80%, ICC 0.6-1, **Figure 2A**), and include exposures that are widespread in the population, and with stable levels in individuals. A large proportion of these substances were PFAS and chlorinated analytes, which have biological persistence and long pharmacokinetic half-lives. For example, PFHxS had an ICC of 0.94 (100% DF, reported half-life 8.5 yrs)(64) and PFOS isomers had ICCs of 0.95 (100% DF, reported half-life 5.4 yrs for linear PFOS)(64). Another example was the fungicide transformation product, chlorothalonil-4-hydroxy (ICC 0.80, 100% DF; here confirmed at Level 1). The half-life of chlorothalonil-4-hydroxy in humans has not yet been determined, but the high ICC reported here and in a previous study of Costa Rican women (ICC 0.81)(44) emphasizes the importance of further studies for this environmentally persistent chemical. Other Type 2 substances nevertheless included analytes with reported fast elimination half-lives in the range of days or hours. Such examples were 3-hydroxycotinine (ICC 0.91, 88% DF, half-life 6.6 hr(65); here confirmed at Level 1), and the pesticide pentachlorophenol (ICC 0.83, 98% DF, half-life 20 d(66); Level 1). These latter results suggest that, despite relatively rapid elimination pharmacokinetics, consistent lifestyle factors or behaviors over the course of two years (e.g. tobacco use) can result in steady levels of environmental substances in blood plasma. In fact, the ICCs for the above environmental chemicals were comparable to that of the targeted steroid hormone, testosterone (ICC 0.95, 100% DF; Level 1). While substances in the Type 2 category do not necessarily require a longitudinal design with repeated sampling to accurately classify exposure in health studies, longitudinal exposome studies with untargeted chemical exposomics may be a powerful approach to discovering emerging persistent chemicals in the human population. For example, among the thousands of non-annotated molecular features detected in this study, these could be prioritized for identification based on simultaneously high ICC and detection frequency.

Type 3 exposures were categorized as common-unstable (detection frequency > 80%, ICC < 0.4, **Figure 2A**). Substances in this category are widespread in the population, but have unstable levels in individuals, thus the health relevance of these can most powerfully be explored in longitudinal studies with repeated measurements of exposure. Examples here included substances associated with exposure through diet, and/or having short elimination half-lives, such as the insecticide metabolite 3,5,6-trichloro-pyridinol (ICC 0.25, 97% DF, half-life 27 hr(67); Level 1), with exposure sources linked to the consumption of imported foods in Sweden.(68) Other examples are the caffeine metabolite theobromine (ICC 0.31, 100% DF, half-life 7-12 hr(69); Level 2), azelaic acid which is found in grains(70) (ICC 0.40, 100% DF; here confirmed at Level 1, **Table 1**, **Figure S19**), and the herbicide ioxynil(71) (ICC 0.37, 92% DF; Level 2). An additional example was the rubber additive 1,3-diphenyl guanidine (ICC <0.01, 83% DF, half-life 9.6 h in rats(23); Level 1), which as discussed above is widespread in the environment.

Finally, Type 4 exposures were categorized as rare-unstable (detection frequency < 20%, ICC < 0.4, **Figure 2A**). These exposures are more difficult to understand with regards to health, and could be deprioritized in exposome studies to further reduce the dimensional complexity and multiple testing. Examples of Type 4 exposures are the industrial chemicals 2,6-di-tert-butyl-4-nitrophenol (ICC 0.33, 20% DF; Level 1) and the isomers 1- and 2-naphthalene sulfonate (ICC 0.29, 13% DF; Level 1), both discussed above.

### Co-Exposures in the Chemical Exposome

The complexity and high dimensionality of the chemical exposome represent major challenges to its implementation in health studies, however it has been proposed that reduction of its dimensional complexity should be possible by grouping correlated exposures.(56) Such co-exposures may arise if two or more substances have common exposure sources and similar pharmacokinetics, resulting in correlated dynamics. Here we highlight several co-exposures among the 46 participants that were evident in HCA heatmaps (**Figure 4A, Figure S22**), and for discussion we group these into ‘common’ and ‘rare’ co-exposures based on whether these were evident among many participants (**Figure 4A** groups **A-F,** zoom-in **Figure S23**), or only in individuals or a small fraction of the population (**Figure 4A** groups **G-N,** zoom-in **Figure S24**), respectively. Both HCA heatmaps are based on all 46 participants and 519 annotated substances (Level 1 and 2), but for simplicity of discussion we focus on **Figure 4A**, which is based on the average response of each substance for each individual’s multiple visits (n=6). **Figure S22** shows the chemical exposome at each visit to simultaneously visualize intra- and inter-individual patterns. To guide interpretation and hypothesis generation, both heatmaps are interactive (available to navigate at https://s3wp-exposomics.serve.scilifelab.se) and metadata are displayed for each substance (i.e. chemical class, subclass, confidence level, ionization mode, DF) and participant (sex, age, BMI).

**Figure 4.**
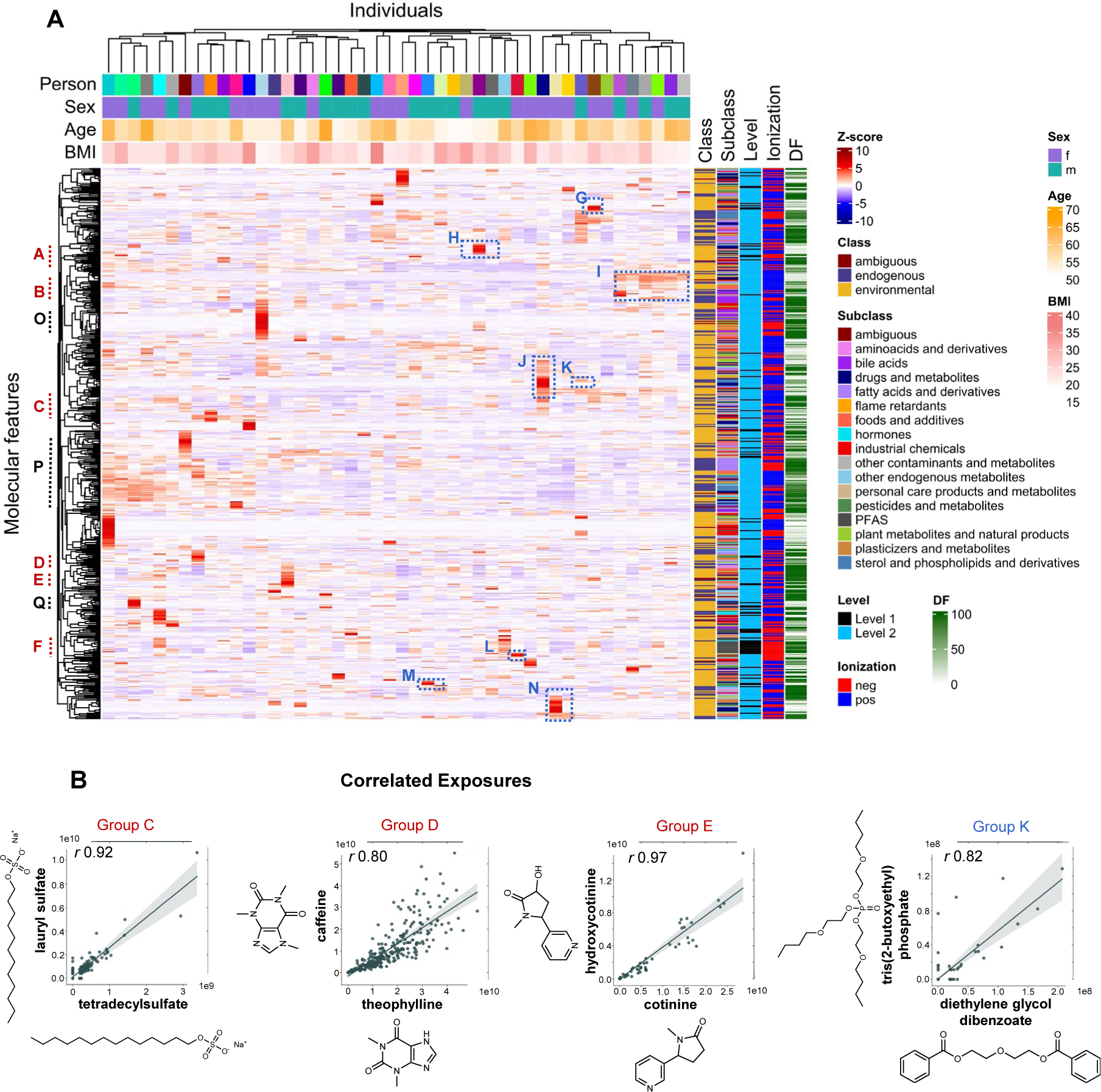
Clustered exposome profiles of S3WP participants and example correlated exposures. In panel (A), a hierarchical cluster analysis heatmap with dendrograms shows the exposome profiles of 46 individuals, each averaged across the 6 clinical visits for 519 annotated substances (Level 1 and Level 2). Color coding of chemical substances (“molecular features”) is according to class, subclass, confidence level of identification, ionization and detection frequency (DF). Color coding of individuals is according to sex, age and BMI. Groups of interest are highlighted on the heatmap (in red A-F: common co-exposures; in blue G-N: rare co-exposures; in black O-Q: endogenous metabolites, see zoomed in versions in Figures S23, S24, S27). Common co-exposures were detected in many participants while rare co-exposures were detected only in individuals or a small population fraction. Panel (B) shows linear regression plots with 95% confidence intervals across all 6 visits (n= 276) between features that clustered together in the heatmap. The Pearson correlation coefficient (*r*) is shown for each regression and all analyte pairs showed significant correlation (p-value < 0.001).

One common co-exposure (Group A, **Figure 4A**, zoom-in **Figure S23**) included the correlation between the targeted fungicide pentachlorophenol (ICC 0.83, 98% DF) and the dietary fatty acid, pentadecanoic acid, which is a biomarker of dairy consumption(72) (Level 2; ICC 0.29, 93% DF; *r* 0.43, p < 0.005). Pentachlorophenol is also a metabolite of other organochlorines, including the pesticide hexachlorobenzene(73) which has been associated with dairy consumption in Sweden.(74) Another common co-exposure (Group B, **Figure S24**) included homologues of polyethyleneglycol (Level 2, ICC 0-0.31, 36-100% DF), which are approved by the US Food and Drug Administration for dermal, oral and intravenous pharmaceutical applications,(75) and have been detected in environmental samples,(76,77) but not in human samples previously. Group C co-exposures (**Figure S23**) included sodium lauryl sulfate (ICC 0.09, 45% DF, confirmed Level 1), strongly correlated with tetradecyl sulfate (Level 2; ICC 0.07, 36% DF; *r* 0.92, p < 0.001, n = 276, **Figure 4B**) and lauryl diethanolamide (Level 2; ICC 0.01, 32% DF; *r* 0.66, p < 0.001, n = 276), all of which are ingredients in household cleaning and personal care products, such as detergents, shampoos, liquid soap and cosmetics.(49,50,78) Correlations such as these, between confirmed analytes (targeted/Level 1) and one or more Level 2 substances in a related chemical class provide further confidence in the Level 2 annotations, and other examples are discussed below.

Another common co-exposure (Group D, **Figure S23**) involved strong correlations among the targeted analyte caffeine (ICC 0.50, 99% DF), and the Level 2 caffeine metabolites(79) theobromine (ICC 0.31, 99.6% DF; *r* 0.60, p < 0.001, n = 276 individual samples) and theophylline (ICC 0.52, 100% DF; *r* 0.80, p < 0.001, n = 276, **Figure 4B**). While these three alkaloids are typical of coffee plants, they also clustered with 4-methylcatechol (Level 2; ICC 0.60, 99% DF), a compound generated during the roasting of coffee beans.(80) Similarly, in Group E (**Figure S23**) the targeted nicotine metabolite, cotinine (ICC 0.94, 20% DF), was strongly correlated with a secondary nicotine metabolite, 3-hydroxycotinine(81) (ICC 0.91, 88% DF; *r* 0.97, p < 0.001, n = 276, **Figure 4B**), initially discovered and annotated at Level 2 but confirmed at Level 1. Group F (**Figure S23**) shows the prominent PFAS class as a co-exposure cluster (i.e. many PFAS were highly correlated, **Figure S25**). Additionally, the HCA heatmap with all 6 visits (**Figure S22**) provides visual evidence for the high ICCs of these substances, whereby the majority of individuals showed consistently high (red) or low (blue) responses across all visits (also see zoom-in subset in **Figure 5A**).

**Figure 5.**
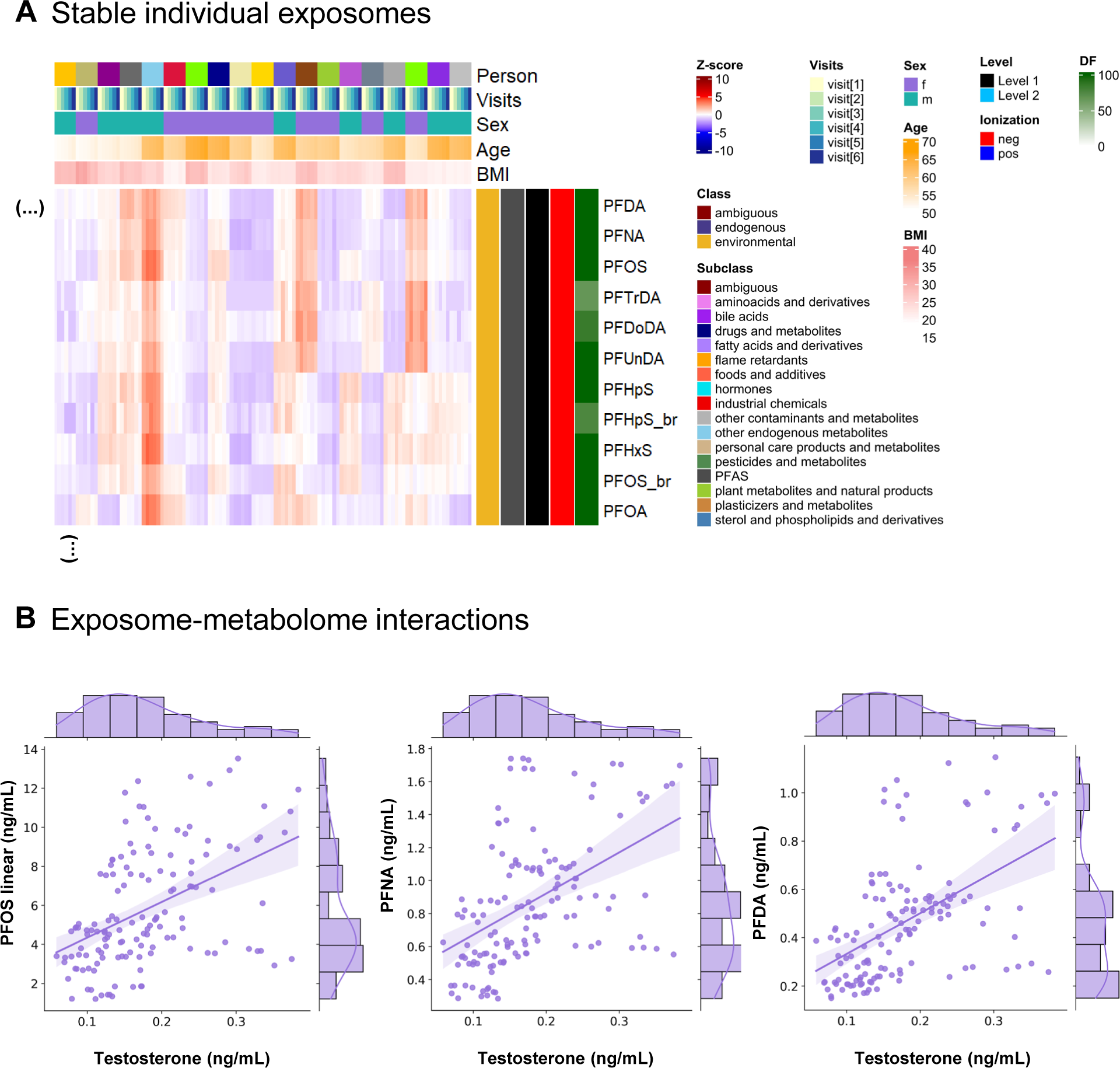
Stable PFAS exposome profiles and example exposome-metabolome interactions. Panel (A) shows a zoomed-in subset of the hierarchical cluster analysis heatmap (19 out of 46 participants) with 6 visits shown per individual where a stable group of PFAS is observed. Color coding of features is according to class, subclass, confidence Level of identification, ionization and detection frequency (DF). Color coding of individuals is according to sex, age and BMI. Panel (B) show linear regression plots with 95% confidence intervals between testosterone and PFAS targets for female individuals across all 6 visits (n= 138). All analyte pairs showed significant correlation (p-value < 0.001). The plots also show the analyte distributions in histograms.

Examples of rare co-exposures involved groups of chemicals occurring at low detection frequency in the population (approx. < 20%), but also include distinctively higher levels of more commonly detected substances in individuals. In Group G (**Figure 4A**), one distinctive individual had multiple rare co-exposures (zoom-in **Figure S24**), including to triphenyl phosphine oxide (ICC 0.64, 3.2% DF; Level 1) which was detected in all 6 plasma samples of this individual. As noted above, this substance is used as an intermediate in pharmaceutical products,(29) thus it was interesting to note correlated co-exposures in this individual to 2 drugs, trimethoprim(82) (Level 2, ICC <0.01, 14% DF) and ketoprofen(83) (Level 2, ICC 0.34, 57% DF), as well as to the targeted flame retardant, bis(1,3-dichloro-2-propyl) phosphate (ICC <0.01, 12% DF) and the long-chain PFAS, perfluorotetradecanoate (PFTeDA; ICC 0.31, 9% DF). The reasons for these shared rare exposures are not currently understood, but deserve follow-up study, and would not have been uncovered without multiclass targeted and untargeted chemical exposomics.

Another rare co-exposure (Group I, **Figure S24**) involved 6 individuals with similarly high relative exposure to the antibiotic drug metabolite clindamycin sulfoxide(84) (Level 2; ICC 0.54, 9% DF), the uremic toxin and food derived compound(85) alpha-hydroxyhippuric acid (Level 2; ICC 0.81, 17% DF), the fatty acid metabolite 5-hydroxyeicosatetraenoic acid(86) (5-HETE, Level 2; ICC 0.44, 22% DF), and two putative fungal metabolites, i.e. beta-orcinol(87) (Level 2; ICC 0.45, 37% DF) and pestalotin(88) (Level 2; ICC 0.24, 10% DF). In Group K (**Figure S24**), two rare exposures, namely the targeted flame-retardant and plasticizer tris(2-butoxyethyl) phosphate (ICC 0.01, 9% DF) and the plasticizer diethylene glycol dibenzoate (Level 2; ICC < 0.01, 9% DF), were strongly correlated (*r* 0.82, p < 0.001, n = 276, **Figure 4B**).

Additionally, in Group L, (**Figure S24**) the targeted personal care product triclosan (ICC 0.45, 16% DF) clustered with several other chlorinated substances, such as the herbicide 4-chlorophenoxyacetic acid(89) (Level 2; ICC 0.50, 48% DF), the pesticide chloroxynil(90) (Level 2; ICC 0.01, 33% DF), and an isomer of chlorophenol (Level 2; ICC 0.16, 2.2% DF), the latter discovered based on exact mass, chlorine isotopic pattern (**Figure S11**) and retention time adjacent to 4-chlorophenol (confirmed Level 1). In Group M, the targeted herbicide diuron (ICC < 0.01, 20% DF) clustered with the targeted fungicide carbendazim (ICC < 0.01, 4% DF) and the herbicide ioxynil(71) (Level 2; ICC 0.37, 92% DF) (**Figure S24**).

### Chemical Exposomics as a Tool for Precision Health

An individual’s drug metabolism capacity is influenced by both genetic and environmental factors, and understanding these factors are key objectives of personalized medicine, for example towards precision dosing of drugs.(91) We hypothesized that untargeted chemical exposomics could be a useful tool for monitoring drug efficacy (or toxicity) at the individual level by monitoring drugs and detecting their known or unknown metabolites in plasma. We first observed that groups of rare co-exposures often consisted of drugs and their known metabolites (**Figure S24**). For instance, the anti-inflammatory drug diclofenac (Level 1; ICC 0.19, 6.9% DF) and 5-hydroxy diclofenac(92) (Level 2; ICC 0.37, 9.8% DF) in Group H, the antihistamine cetirizine(93) (Level 2; ICC 0.88, 21% DF) and hydroxyzine (Level 2; ICC0.83, 68% DF) in Group J, and the antidepressant sertraline (Level 2; ICC 0.32, 2.9% DF) and norsertraline(94) (Level 2; ICC 0.33, 2.5% DF) in Group N. To further explore the potential of chemical exposomics to identify drug metabolites whose MS2 spectra are unknown, or not publicly available, we suspect-screened all 38499 non-annotated untargeted features detected in ESI-for: (i) theoretical m/z corresponding to hydroxyl, sulfate or glucuronide metabolites of the above mentioned pairs of parent drugs and metabolites, and (ii) correlation with the respective parent drug or metabolites (i.e. p < 0.001, n = 276 individual samples, **Figure S26**). By these combined criteria, a suspected additional hydroxy metabolite of diclofenac (m/z 310.0049, RT 10.74 min, 2 ppm error, ESI-) was found that correlated both with diclofenac (*r* = 0.61, m/z 294.0094, RT 13.16 min, ESI-) and 5-hydroxy diclofenac (*r* = 0.92, m/z 312.0190, RT 11.42 min, ESI+), and the corresponding suspect sulfate conjugate (m/z 389.9616, RT 9.50 min, 1.3 ppm error, ESI-) correlated moderately with diclofenac (*r* = 0.63), and strongly with the new hydroxy metabolite (*r* = 0.98). Finally, a suspected glucuronide of sertraline (sertraline carbamoyl-O-glucuronide, m/z 524.0884, RT 13.75 min, 0.1 ppm error, ESI-) correlated strongly with both sertraline (*r* = 0.97, m/z 306.0812, RT 14.09 min, ESI+) and norsertraline (*r* = 0.98, m/z 292.0653, RT 14.31 min, ESI+). These results demonstrate that untargeted chemical exposomics is a feasible tool to inform the pharmaceutical treatment of individuals.

Hierarchical clustering of natural compounds and endogenous metabolites (Level 2, **Figure 4A** groups **O-Q,** zoom-in **Figure S27**) was also observed, including for bile acids (group O), lipids (group P) and microbial metabolites (group Q). These may be useful indicators of behaviour (e.g. dietary habits), or biomarkers of disorders or disease risk. For example, In group O, a single individual deviated from the rest of the population due to high relative response of several bile acids, including cholic acid, glycochenodeoxycholic acid and ursocholic acid (**Figures S27 and S28A**), which can indicate hepatic impairment.(95) Moreover, 12 participants clustered with relatively high responses of several lipids, including the fatty acids cis-4,10,13,16-docosatetraenoic acid (ICC 0.55, 100% DF), 9Z, 12Z-linoleic acid (ICC 0.44, 100% DF) and arachidonic acid (ICC 0.42, 100% DF), with high correlations among these (Group P, **Figures S27 and S28B**). High correlations were observed between the microbial metabolite phenylacetylglutamine (ICC 0.65, 100% DF) and the two uremic toxins p-cresol sulfate (ICC 0.62, 100% DF; *r* 0.80, p < 0.001, n = 276) and 3-indoxyl sulfate (ICC 0.51, 100% DF; *r* 0.49, p < 0.001, n = 276) (Group Q, **Figures S27 and S28C**). This microbial metabolite and the associated uremic solutes accumulate in patients with chronic kidney disease, and their levels have been associated with adverse outcomes including cardiovascular disease.(18,96)

### Exposome-Metabolome Interaction

Correlations between environmental chemicals and endogenous metabolites can be suggestive of exposome-metabolome interactions, whereby the exposure(s) induce a metabolic response. Although the current study is relatively small, we examined for evidence of endocrine disruption between testosterone and PFAS, which are quantitative targeted analytes that were detectable in most samples. Testosterone and PFAS did not cluster together in HCA heatmaps because men have much higher testosterone levels than women, and any correlations may be sex-specific. Among all females, positive correlations were observed between testosterone and several PFAS (**Figure 5B**), including linear PFOS, perfluorononanoate (PFNA) and perfluorodecanoate (PFDA). These three associations were statistically significant in mixed-effects models, which account for the repeated sampling of individuals, and PFOS and PFDA remained significant in adjusted models controlling for baseline age and BMI (**Table S8**; p<0.02 after Bonferroni correction). Positive associations between PFOS, PFOA and PFHxS with total and free testosterone have been reported in postmenopausal women (median 63 yrs),(97) similar to the age of women in the current study (median 57 yrs). However, the same study reported no similar effect for PFNA and PFDA on any reproductive hormones.(97) These findings for PFAS deserve further study considering that a meta-analysis of prospective studies has reported that higher blood testosterone was associated with increased risk of breast cancer in postmenopausal women.(98)

### Advances, Limitations and Future Directions

The current study represents the first application of longitudinal chemical exposome profiling in human blood of multiple individuals, and therefore provides new tools and valuable context for designing and executing larger statistically powered studies of disease etiology. In terms of tools, the combined targeted and untargeted chemical exposomics workflow showed promise by enabling sensitive and precise quantification of priority targeted analytes (e.g. pollutants and hormones) with parallel potential to discover molecules of environmental and health-relevance. As a result, in this longitudinal study of only 46 individuals we reported significant time-trends, exposome-metabolome interactions, and novel co-exposures to natural and man-made substances, aided by visualizing exposomes in interactive heatmaps from hierarchical clustering. These results generally show promise for chemical exposomics in precision health and precision medicine applications.(99) Nevertheless, the vast majority of molecular features detected in the untargeted analysis remain unidentified ‘molecular dark matter’, and the underlying laboratory methods and data analyses must still be scaled up to identify risk factors of disease in larger studies.

It has been envisioned that unravelling the effects of the chemical exposome on progression of disease will require network science and systems biology approaches to integrate chemical and biomolecular interactions across many levels of cellular organization.(56) Thus, from a molecular epidemiology perspective, the range of mean ICCs reported here from parallel measures of the chemical exposome, metabolome, proteome, lipidome and microbiome provide important context and demonstrate the practical temporal challenge to this network approach. More specifically, although each omic tool is accurate and reproducible in its own right, the representativeness of the resulting molecular profiles is lowest for the exposome because it changes more dynamically over time. The wide range of ICCs for hundreds of endogenous and exogenous small molecules reported here can therefore assist in design (i.e. sample sizes and sampling frequencies) of longitudinal cohorts focused on health impacts of the chemical exposome.

## Materials and Methods

### Recruitment and Sampling

Plasma samples were from healthy participants enrolled in the Swedish SciLifeLab SCAPIS Wellness Profiling (S3WP) study(57) whom were previously recruited to the Swedish CArdioPulmonary bioImage Study (SCAPIS), a larger cohort of 30,000 participants aged 50-65 years and representative of the general Swedish population.(57,100) The subjects were extensively phenotyped before entering the S3WP study through questionnaires and examinations for clinical chemistry, anthropometry and imaging to assess coronary and carotid atherosclerosis.(57,100) The S3WP study included 6 examination visits in two rounds, with 94 of 101 enrolled subjects completing all 6 visits.(57) In the first round, an examination was performed every third month (±2 weeks), resulting in four visits per participant. In the second round, an examination was performed approximately at 6-month intervals, resulting in two additional visits per participant. At each visit, samples of blood, urine and feces were collected.(57) All subjects fasted overnight before the visits.(57) In the present work, longitudinal sample sets of plasma (50 - 200 µL, 6 samples per person, 276 total samples) collected between the end of 2015 and start of 2018, were randomly selected from 46 of the S3WP participants, so that 23 males and 23 females were included, with birth years in the ranges 1950-1955 (n=14), 1956-1960 (n=16) and 1961-1965 (n=16). The study was approved by the Ethical Review Board of Göteborg, Sweden, and all participants provided written informed consent. The study protocol conforms to the ethical guidelines of the 1975 Declaration of Helsinki.

### Sample Preparation

Plasma samples were prepared and analyzed following a combined targeted and untargeted chemical exposomics method described by Sdougkou et al.(7) Briefly, 200 µL plasma aliquots were placed in 2 mL tubes and fortified with 10 µL of isotopically-labelled internal standard mixture (34 substances) in methanol (MeOH; Optima LC/MS Grade, Thermo Scientific) (**Table S1**, final concentration 1 ng/mL of each). Protein precipitation was by adding 800 µL acetonitrile (Optima LC/MS Grade, Fisher Chemical) containing 0.5% citric acid (CA) (BioUltra, anhydrous, ≥ 99.5%), vortexing for 20 s at 4 °C for 20 min, and centrifugation (20,800 x *g*) at 4 °C for 10 min. Supernatants were loaded to HybridSPE-Phospholipid cartridges (500 mg/6 mL, Merck) prewashed with 12 mL MeOH and 12 mL ACN containing 0.5% CA. Elution was with 1 mL ACN containing 0.5% CA, followed by 2 mL MeOH containing 1% ammonium formate (LiChropur, ≥ 99.0%) into 15 mL tubes. The pH of the extracts was adjusted from approximately 3 to 6.5 by adding 40 µL of 25% ammonia solution (LiChropur, LC-MS grade), and centrifuging for 10 min at 4,300 x *g* at room temperature. Supernatants were transferred to 5 mL tubes, evaporated to 100 µL under nitrogen flow and ultrasonicated for 5 min. A final rinse of the tubes with 100 µL MeOH was performed to reach a final extract volume of 200 µL, followed by centrifugal filtration (10,600 x *g*) for 10 min (0.2 μm nylon filters, Thermo Scientific). The final extracts were transferred to amber glass vials and spiked with 10 μL of diuron-d6 solution (final concentration 4 ng/mL) to correct for extract volume variations and to monitor instrumental performance.

### LC-HRMS Analysis

Measurements were conducted by ultrahigh pressure LC (Ultimate 3000, Thermo Scientific) with HRMS acquisition (Q Exactive Orbitrap HF-X, Thermo Scientific) in positive and negative electrospray ionization mode (ESI+ and ESI-) as previously described.(7) Spectral acquisition alternated between full scan (i.e. MS1; 90-1000 mass-to-charge ratio (m/z), 120,000 nominal resolution) and data-independent MS/MS acquisition (DIA) (i.e. MS2; 30,000 nominal resolution, product ion scan range starting from 50 Da) with four *m/z* precursor windows of equal size (237 Da), with each window overlapping by 10 Da. Data-dependent acquisition (DDA) with an inclusion list of precursor ions was used for analyte confirmations. Injection volumes were 20 μL, corresponding to 20 µL plasma-equivalents on-column, and chromatography was at 40 °C on an Acquity BEH C18 column (130 Å, 1.7 μm, 3 x 100 mm, Waters) with an Acquity BEH C18 1.7 µM vanguard pre-column. Upstream of the injector, an Acquity BEH C18 column (130 Å, 1.7 μm, 3 x 30 mm, Waters) was placed to separate instrumental background from sample analytes. A binary gradient elution at 0.4 mL/min used mobile phases (A) water (Optima LC/MS Grade, Thermo Scientific) containing 1 mM ammonium fluoride(101) (Honeywell Fluka, ≥ 98.0%), and (B) 100% MeOH. The elution gradient started at 5% B, linearly increased to 100% B by 15 min, held until 22 min, returning to initial conditions with 4 min equilibration.

### Targeted Analyte Quantification

The chemical exposomics method was quantitatively validated for 77 targeted analytes, including environmental contaminants, dietary chemicals, tobacco markers, drugs and endogenous steroid hormones(7) (method limits of quantification (MLOQ), **Table S2**). Four per- and polyfluoroalkyl substances (PFAS) precursors (perfluorooctane sulfonamidoacetic acid (FOSAA), as well as its N-methyl and N-ethyl derivatives (NMeFOSAA and NEtFOSAA) and N-methyl perfluorobutane sulfonamide (NMeFBSA)), and the pesticides fenuron and carbendazim were added to the method after preliminary validation, bringing the total of the targeted analytes to 83 (instrumental LOQ, **Table S2**). For three target PFAS analytes (perfluorooctane sulfonate (PFOS), perfluorohexane sulfonate (PFHxS), and perfluoroheptane sulfonate (PFHpS)) when branched isomers were detected in plasma these were quantified separately from the corresponding linear isomers.

Peak areas were integrated using Xcalibur Quan Browser (Thermo Scientific, v.4.1), and solvent-based calibration curves with internal standards (9 points, 0.01-100 ng/mL) were used to quantify most targeted analytes. The 276 plasma samples were extracted and analyzed in eight batches, and in each batch the calibration curves were run 3 times (beginning, middle, end). Pooled Swedish reference plasma from a separate cohort was also run multiple times in each injection sequence to support reference standardization(21) quantification of steroid hormones, and to enable retrospective semi-quantification of discovered substances in the untargeted analysis. To support reference standardization, standard addition curves were employed to quantify the discovered substances in the pooled Swedish reference sample. For data summaries and statistics, when analytes were detected at concentrations lower than the respective MLOQ, the analyte concentrations were substituted by MLOQ/2, and when analytes were non-detect, the concentrations were substituted by MLOQ/4.

### Untargeted Data Processing and Structural Annotations

For untargeted analysis and spectral library matching, raw data were processed in MS-DIAL (v.4.90)(102) for feature alignment across samples, MS1 and DIA MS2 spectral deconvolution, and chromatographic peak area integration (parameters in **Table S3**). Each resultant molecular feature was defined by a chromatographic retention time (RT), an MS1 *m/z*, and a deconvoluted MS2 spectrum. For feature annotation, spectral matching was considered with total identification score of > 700 (i.e., corresponding to at least one of dot- and reverse dot-product scores > 600) to MassBankEU (https://massbank.eu/), MassBank of North America (MoNA; https://mona.fiehnlab.ucdavis.edu/) and Global Natural Product Social Molecular Networking (GNPS; https://gnps.ucsd.edu/). For each annotated feature, a class (endogenous / environmental) and subclass (e.g. bile acids, PFAS) was assigned following searches of PubChem,(103) Human Metabolome Database (HMDB)(104) and Food Metabolome Database (FooDB).(105)

The untargeted MS-DIAL feature lists from ESI+ and ESI-were combined and analyzed in Python (v. 3.8.16)^32^ using Jupyter Notebook (v. 6.5.2)^33^. Redundant features from ESI+ and ESI-were identified based on a mass tolerance of 0.002 Da (based on calculated M from [M-H]^-^ in negative, [M+H]^+^ in positive mode) and RT tolerance of 0.2 min, and the corresponding feature with lowest average peak area was discarded. Peak areas of the final combined dataset were corrected for instrumental and batch variation using the principal component analysis (PCA) scores of isotope-labelled internal standard signal intensities in each sample,(106,107) and then normalized by the volume of each plasma sample analyzed.

In MS-DIAL only features with peak areas 5 times higher than the corresponding procedural blanks (2 blanks per batch) were kept (i.e. sample maximum / blank average > 5). In further processing, the ratio of each sample’s peak area to the average procedural blank peak area was computed for each feature; 0.1 was added to the average procedural blank area of each feature to avoid 0 in the denominator when there was no blank response. For each feature, only samples with peak areas at least 5 times higher than the corresponding blank average were considered; areas in each sample that did not pass this threshold were set to 0. Average areas in blank (when detectable) were subtracted from peak areas in samples which passed the threshold. This additional blank filtering / subtraction step constrained potential artefacts in the final dataset that would lead to a misleading higher detection frequency (DF). As a final data reduction step, highly correlating annotated features (Pearson correlation coefficient (*r*) > 0.95, p-value < 0.001) with RTs differing by a maximum of 0.1 min, and with identical peak shapes (assessed by visual inspection) were considered as originating from the same analyte (e.g. produced by in-source fragmentation, or an adduct); the feature with lowest average peak area was discarded.

### Statistical Analysis and Visualization

PCA was performed in SIMCA (v. 17.0, Umetrics) for the targeted dataset. The package ComplexHeatmap (2.14.0)(108) was used to visualize the combined targeted and annotated untargeted dataset in R (v. 4.2.1) and R Studio (v. 2023.03.1). Hierachical cluster analysis (HCA) dendrograms and heatmaps were generated based on Pearson correlation as clustering distance and average linkage as clustering similarity method. For other data visualizations, the Python libraries Plotly (v.5.13.0)(109) and Seaborn (v.0.12.2)(110) were used. The package SciPy (v. 1.7.3)(111) was used for Student’s t-test and one-way ANOVA as well as calculating Pearson correlation coefficients. The package statsmodels (v. 0.13.5)(112) was used for Bonferroni corrections and Tukey’s test. The package Pingouin (v. 0.5.3)(113) was used for the ICC calculations, and the package umap-learn (v. 0.5.3) was used for uniform manifold approximation and projection (UMAP).(114) For UMAP analyses the default parameters were always used: number of neighbors was 15, minimum distance was 0.1 and Euclidean distance was used as a metric. Unit-variance scaling was applied before UMAP or PCA analyses. Mixed-effects linear regression analyses were conducted using the lme4 package (v 1.1-33) in R (v 4.3.0) to examine (a) trends over time for pentachlorophenol and 3,5,6-trichloro-pyridinol, and (b) associations between PFAS and testosterone. All models included participant-specific random effects with random intercepts and slopes. To examine trends over time, concentrations of each analyte (dependent variables) were natural log (ln) transformed, days from baseline (independent variable) were scaled to year, and resulting slopes were first-order rate constants (yr^-1^) used to calculate half-lives. Associations between log_10_ transformed testosterone (dependent variable) and log_10_ transformed PFAS analyte (independent variables) concentrations were examined in unadjusted models, as well as in models adjusted for both baseline age and baseline body mass index (BMI) as fixed-effects, and in both cases the raw p-values were Bonferroni corrected for multiple hypotheses using the ‘p.adjust’ function in R.

## Acknowledgments

None.

## Author contributions

JWM, FE., LF, MU and GB contributed to study design and the grant application to Formas. JWM coordinated the research and contributed to data interpretation and writing. KS contributed to study design, performed HRMS analyses, and with the help of SP performed sample extractions. KS and SP performed data analyses, interpreted the results and created the figures. BB performed confirmations with reference standards. LJMM conducted statistics with mixed-effects models. GB coordinated sample distribution for analysis. KS drafted the main paper with SP and JWM. All authors commented or edited the final version.

## Data and materials availability

All data needed to evaluate the conclusions in the paper are available in the main text or the supplementary materials. Interactive hierarchical cluster analysis heatmaps are available at https://s3wp-exposomics.serve.scilifelab.se. Access to the MS data and participant-level datasets is available upon reasonable request by contacting the corresponding author.

