## Supplementary text, figures, tables for "Longitudinal Exposomics in a Multiomic Wellness Cohort Reveals Distinctive and Dynamic Environmental Chemical Mixtures in Blood"

Kalliroi Sdougkou *et al.*

**This PDF file includes:**

Supplementary Text

Figs. S1 to S29

Tables S6 and S8

References (143 to 145)

**Other Supplementary Materials for this manuscript include the following:**

Tables S1 to S5 and S7 (separate file; XSLX)

Supplementary Text

Quality Control Notes. Samples from the same individuals were analyzed within the same batch, but were randomized in the injection sequence. A targeted analyte that was detected with high detection frequency (55%) but excluded from further data processing and visualizations was bisphenol A (BPA) (median concentration in samples 2 ng/mL, max 401.5 ng/mL). BPA in humans and animals is rapidly metabolized to a glucuronide of BPA, thus only low levels are expected in blood of healthy individuals. (143) Since no field blank was available (representing sampling and storage of samples prior being received), it was considered appropriate to discard BPA quantifications, despite being well above the blank contamination level. An additional proof for discarding this analyte was its strong correlation with the Level 2 annotated feature bisphenol A- (2,3-dihydroxypropyl)-glycidyl ether (BADGE-H2O) (Pearson coefficient 0.92, p-value < 0.001, **Figure S29**). BADGE-H2O is a hydrolysis product of bisphenol A diglycidyl ether (BADGE), which is often used as a monomer in the production of epoxy-based polymers, as well as an additive for the elimination of surplus hydrochloric acid in polyvinyl chloride (PVC) organosol production. (144) BADGE can transform to BADGE-H2O in the environment, (144) and similar to BADGE, BADGE-H2O is unstable in biological matrices. (145)

Intraclass Correlation Coefficient Calculation. To calculate the intraclass correlation coefficient (ICC) for molecular profiles reported by Tebani et al., (57) only individuals present in the exposomics study reported here (n=46) were included. Since, not all 46 individuals and /or all visits were included in each molecular profile in Tebani et al., (57) deviating numbers of samples were included here for the ICC calculation for each molecular profile. More specifically, 22 individuals (4 visits per individual) were used for the lipidome, 46 individuals (4 visits per individual) were used for the metabolome, 44 individuals (6 visits per individual) were used for the proteome and 44 individuals (4 visits per individual) were used for the microbiome ICC calculation.

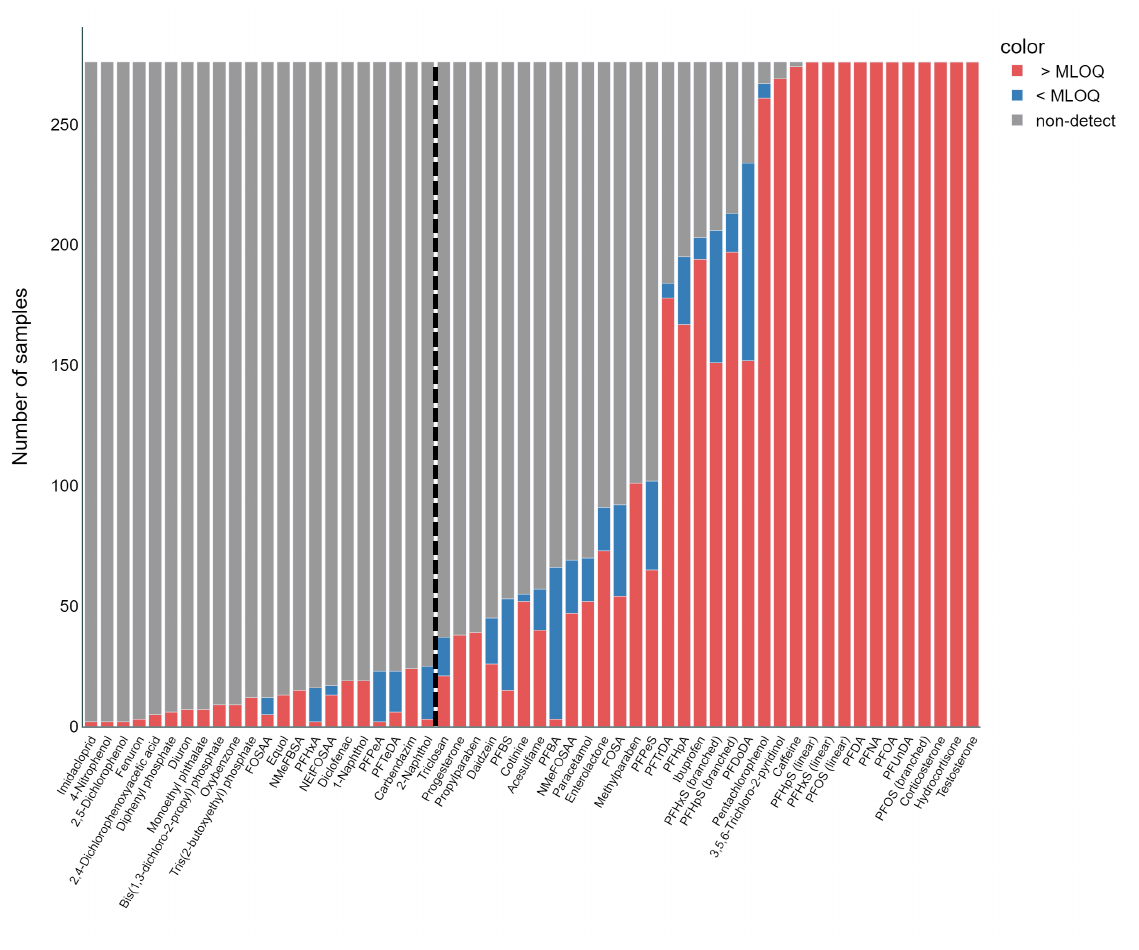

**Figure S1.** Detected target analytes ordered by detection frequency from left to right (1-100%, respectively) among all 276 S3WP plasma samples. Detections above the MLOQ are shown in red, detections below MLOQ are shown in blue, and non-detects are shown in gray. All samples above detection limits (blue and red) were counted towards the detection frequency. The dotted line separates analytes with detection frequency > 10%. For statistical analysis, detections below MLOQ (blue) were substituted by MLOQ/2, and non-detects (gray) were substituted by MLOQ/4.

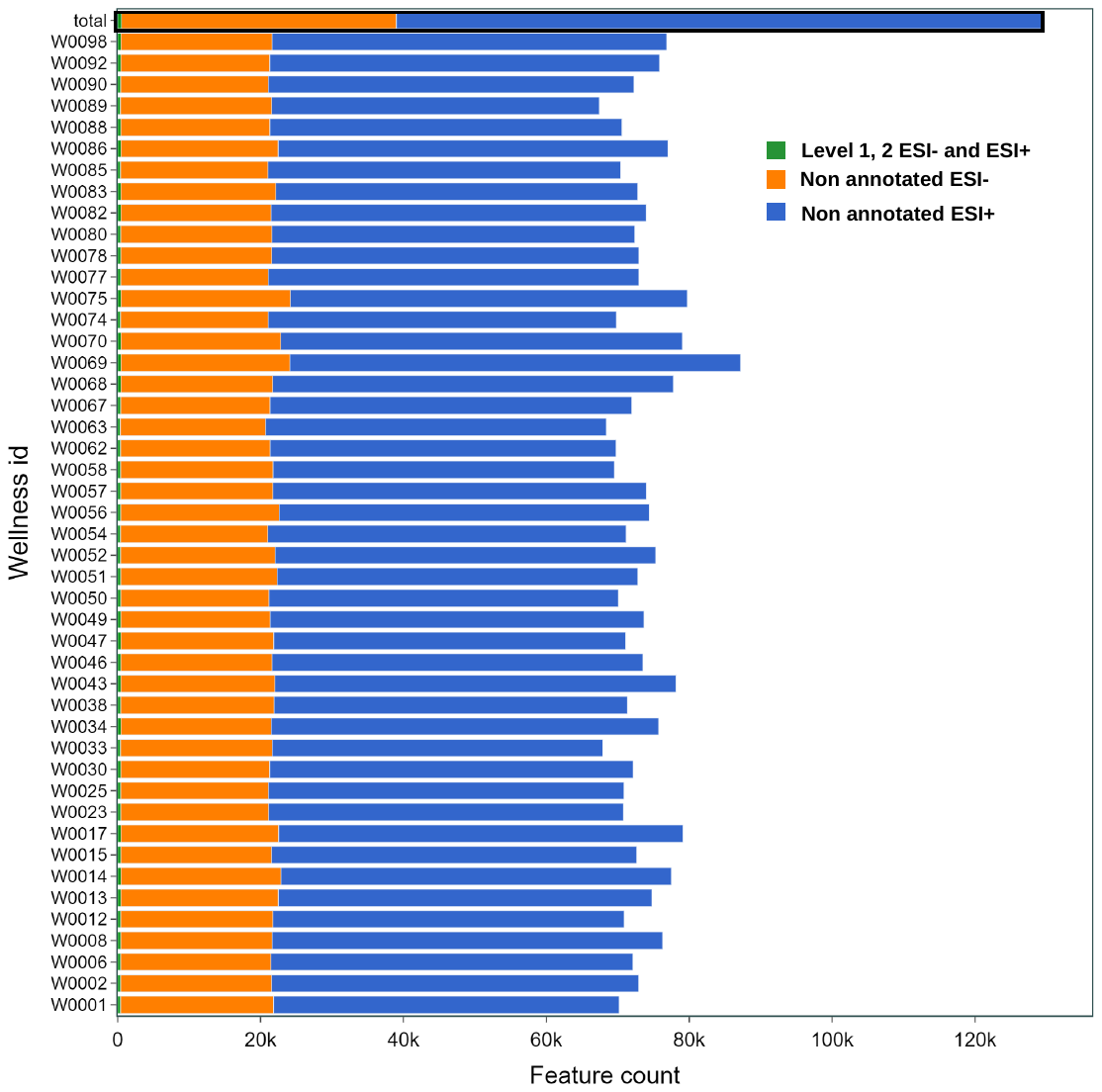

**Figure S2.** Number of annotated features (green, ESI- and ESI+ Level 1 and Level 2) and non-annotated features detected in ESI- (orange) and ESI+ (blue) by untargeted analysis. The feature counts in each category for each individual are a cumulative total number of features detected among all visits, and the top bar (“total”) indicates the cumulative total number of features detected among all individuals and all visits.

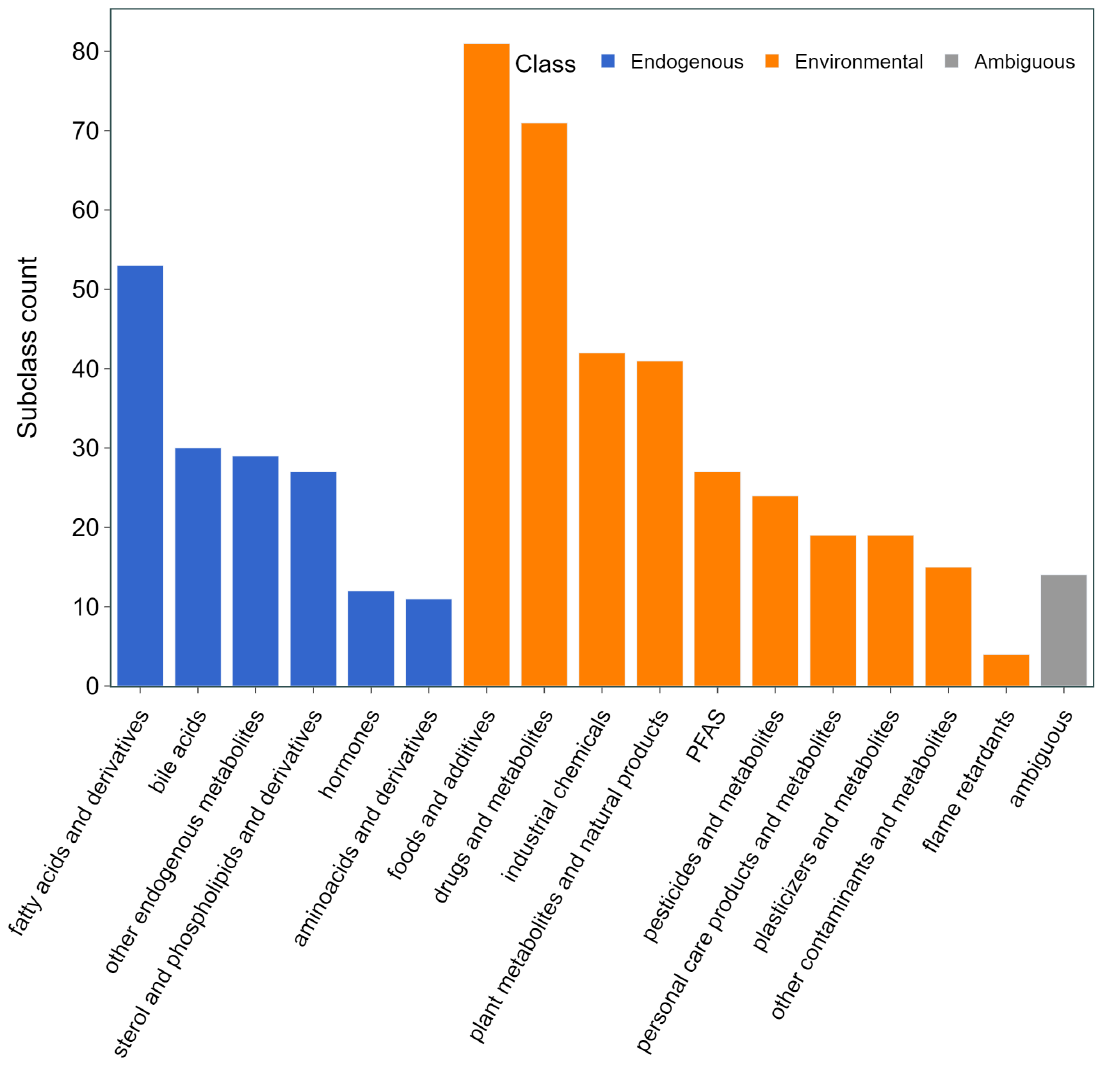

**Figure S3.** Total number of targeted and annotated untargeted analytes by chemical class (blue = endogenous, orange = environmental, gray = ambiguous) and various subclasses (x-axis labels)

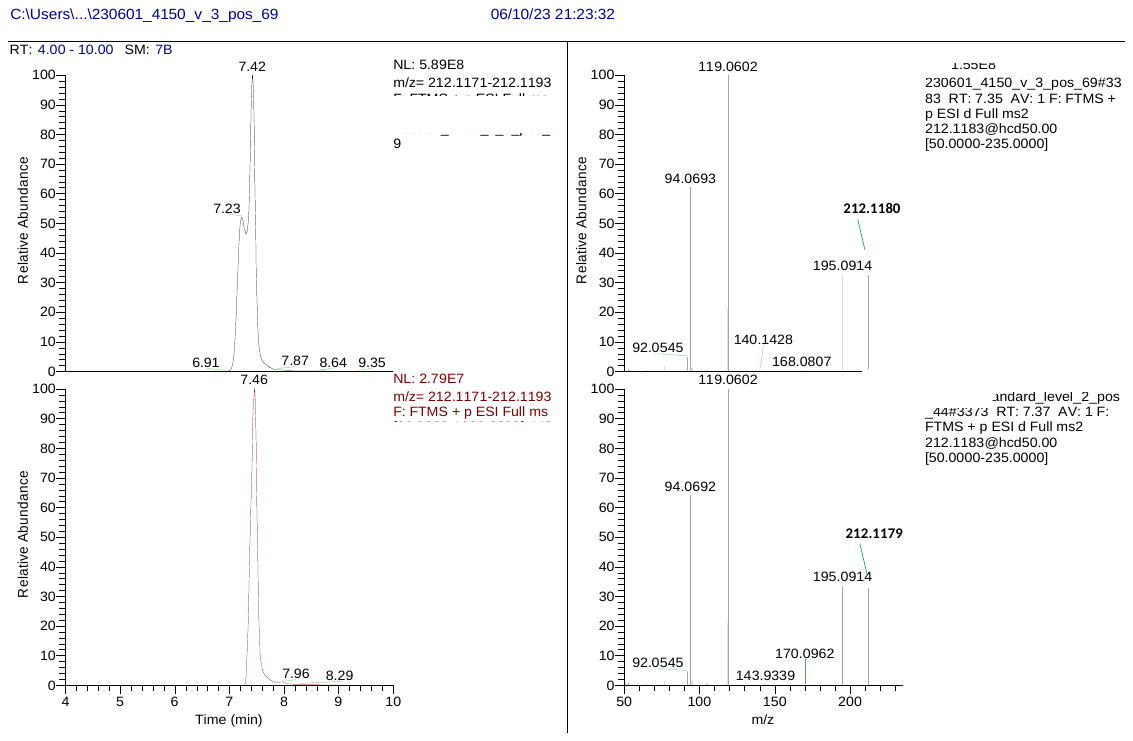

**Figure S4.** Extracted ion chromatogram (left) and data-dependent acquisition (DDA) spectrum (right) for 1,3-diphenylguanidine in individual plasma (top) and standard solution (bottom).

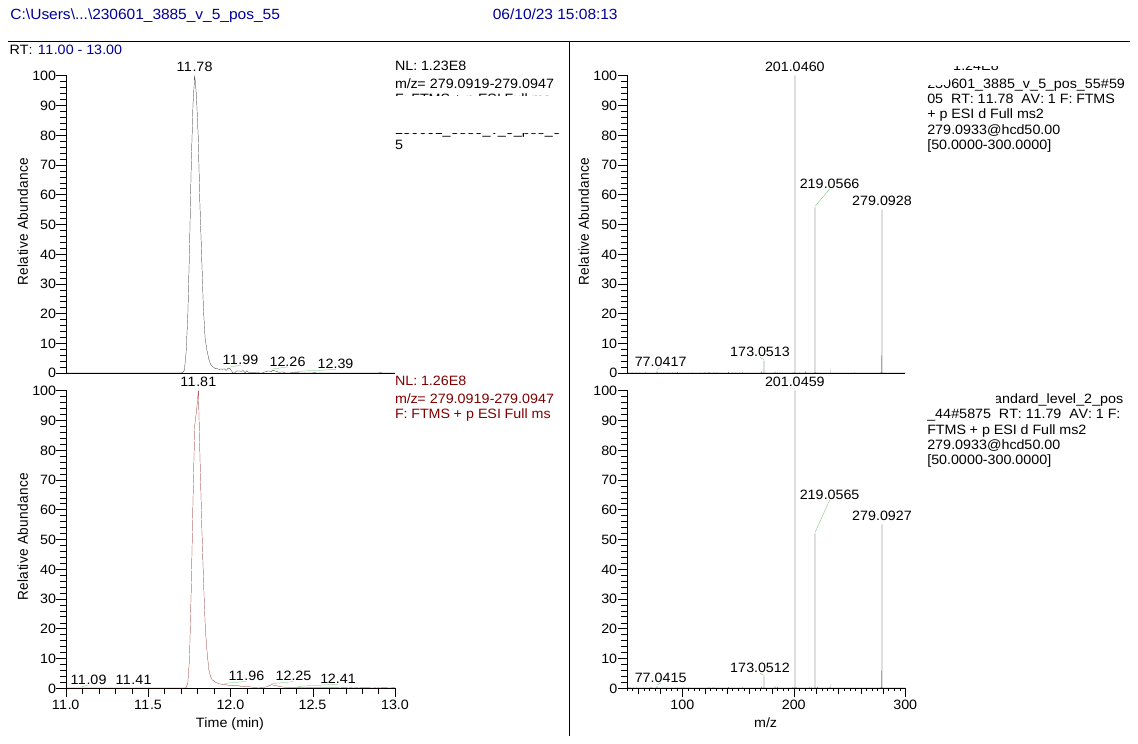

**Figure S5.** Extracted ion chromatogram (left) and data-dependent acquisition (DDA) spectrum (right) for triphenyl phosphine oxide in individual plasma (top) and standard solution (bottom).

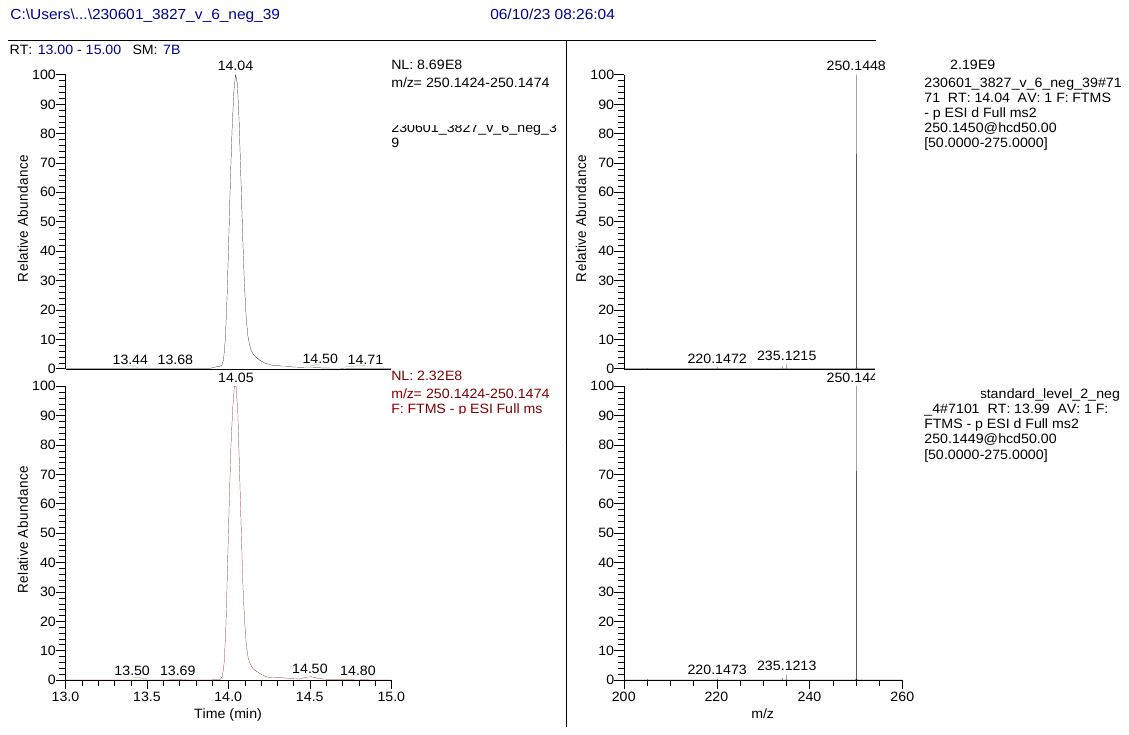

**Figure S6.** Extracted ion chromatogram (left) and data-dependent acquisition (DDA) spectrum (right) for 2,6-di-tert-butyl-4-nitrophenol in individual plasma (top) and spiked pooled plasma (bottom).

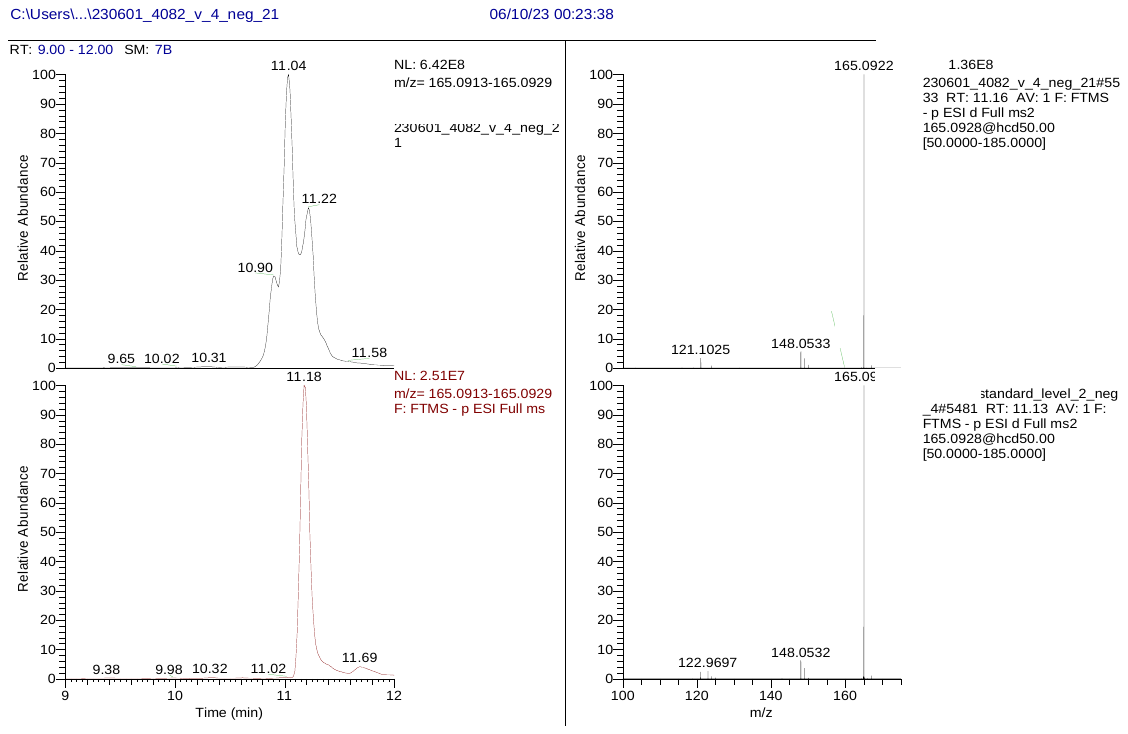

**Figure S7.** Extracted ion chromatogram (left) and data-dependent acquisition (DDA) spectrum (right) for 4-tert-butylpyrocatechol in individual plasma (top) and standard solution (bottom).

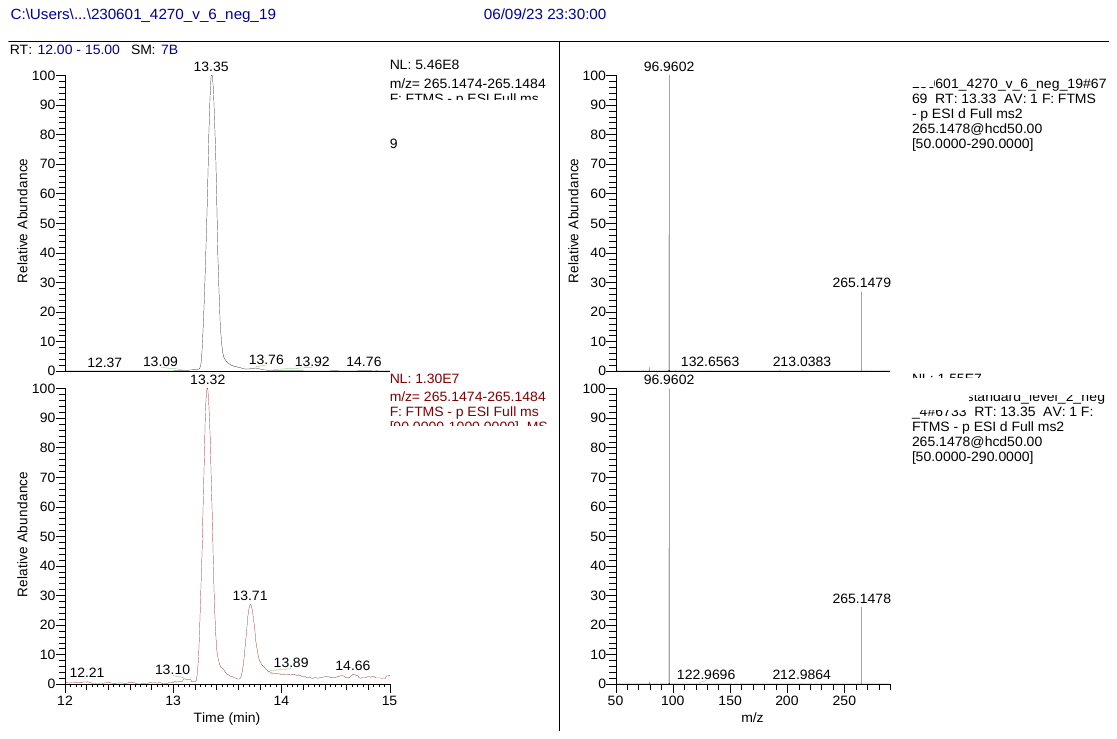

**Figure S8.** Extracted ion chromatogram (left) and data-dependent acquisition (DDA) spectrum (right) for sodium lauryl sulphate in individual plasma (top) and standard solution (bottom). A background peak elutes at 13.71 in the standard solution and 13.76 in individual plasma.

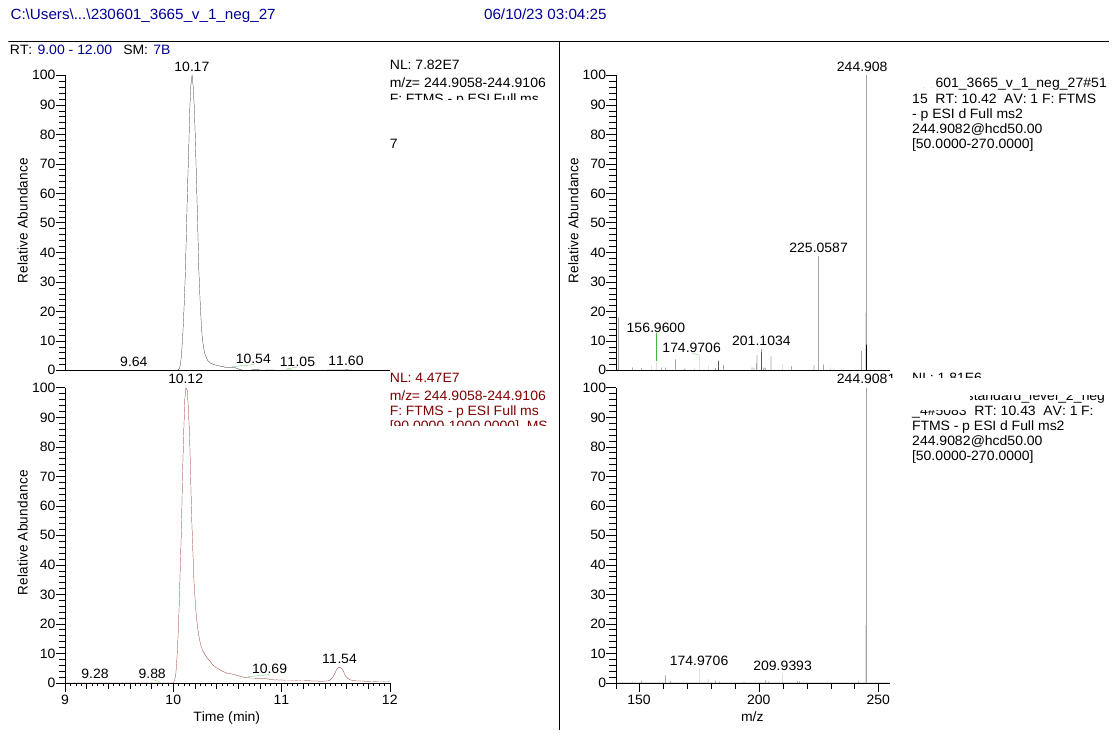

**Figure S9.** Extracted ion chromatogram (left) and data-dependent acquisition (DDA) spectrum (right) for chlorothalonil-4-hydroxy in individual plasma (top) and standard solution (bottom).

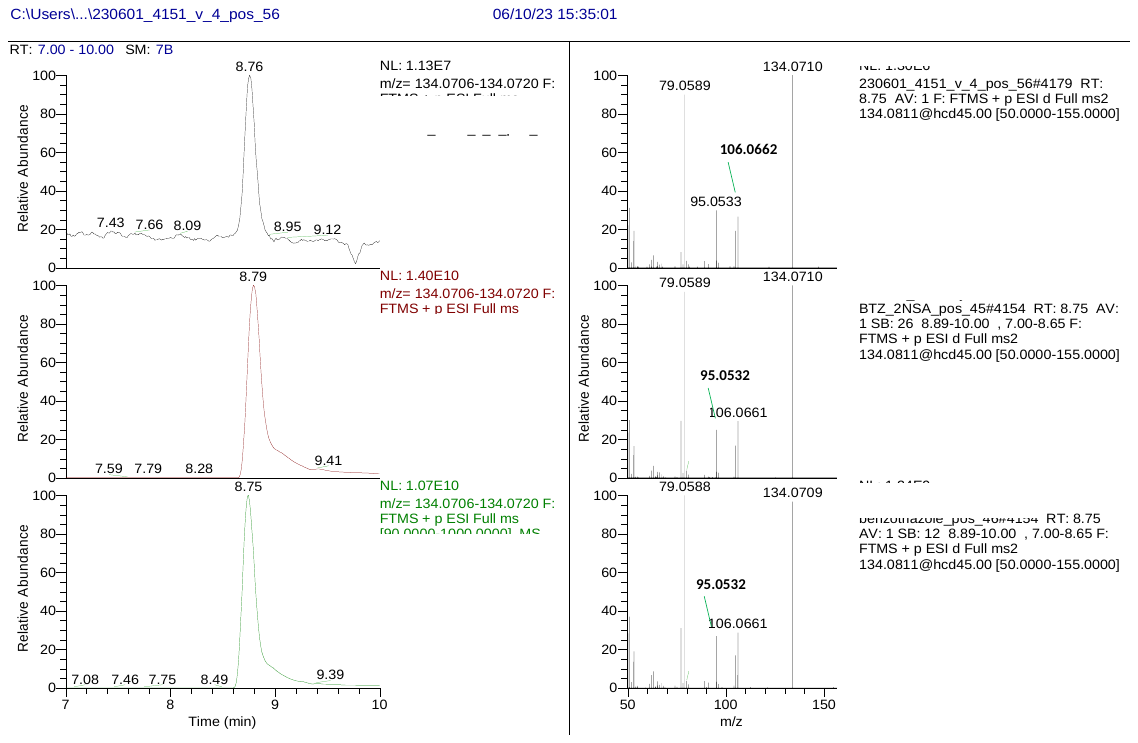

**Figure S10.** Extracted ion chromatogram (left) and data-dependent acquisition (DDA) spectrum (right) for the sum of 4- and 5-methyl-1H-benzotriazole in individual plasma (top) and standard solution (middle: 4-methyl-1H-benzotriazole, bottom: 5-methyl-1H-benzotriazole).

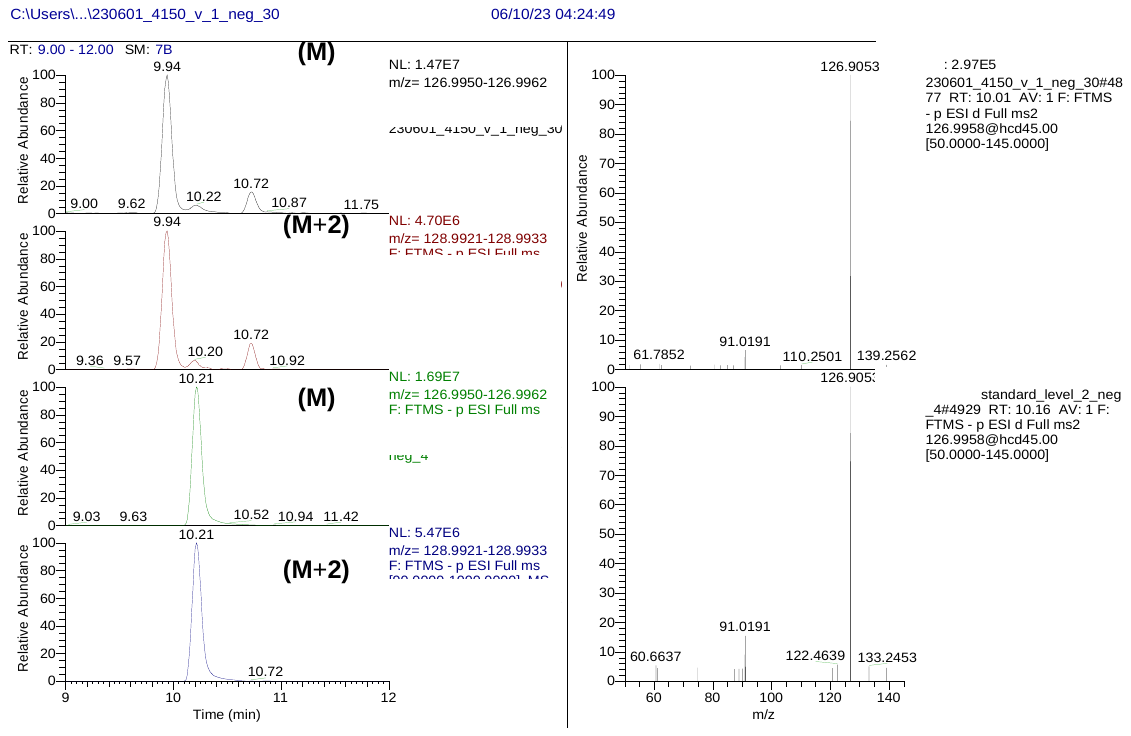

**Figure S11.** Extracted ion chromatogram (EIC) (left) and data-dependent acquisition (DDA) spectrum (right) for 4-chlorophenol in individual plasma (top two EICs and top spectrum) and standard solution (bottom two EICs and bottom spectrum). The EICs are shown in both sample and standard for the M and M+2 ion, corresponding to the chlorine isotopes with atomic masses of 35 Da and 37 Da. 4-chlorophenol elutes at 10.20 min, while chlorinated isomers elute at 9.94 and 10.72 min in the individual plasma.

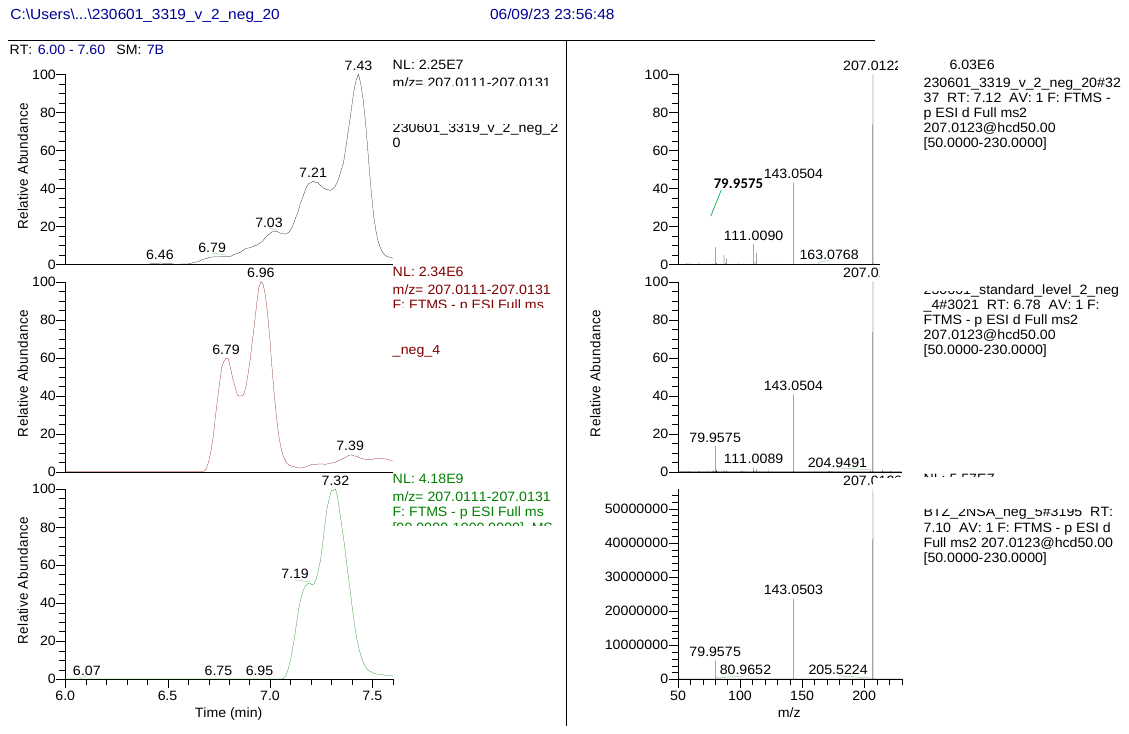

**Figure S12.** Extracted ion chromatogram (left) and data-dependent acquisition (DDA) spectrum (right) for the sum of 1- and 2- naphthalenesulfonate in individual plasma (top) and standard solution (middle: 1- naphthalenesulfonate acid, bottom: 2- naphthalenesulfonate acid).

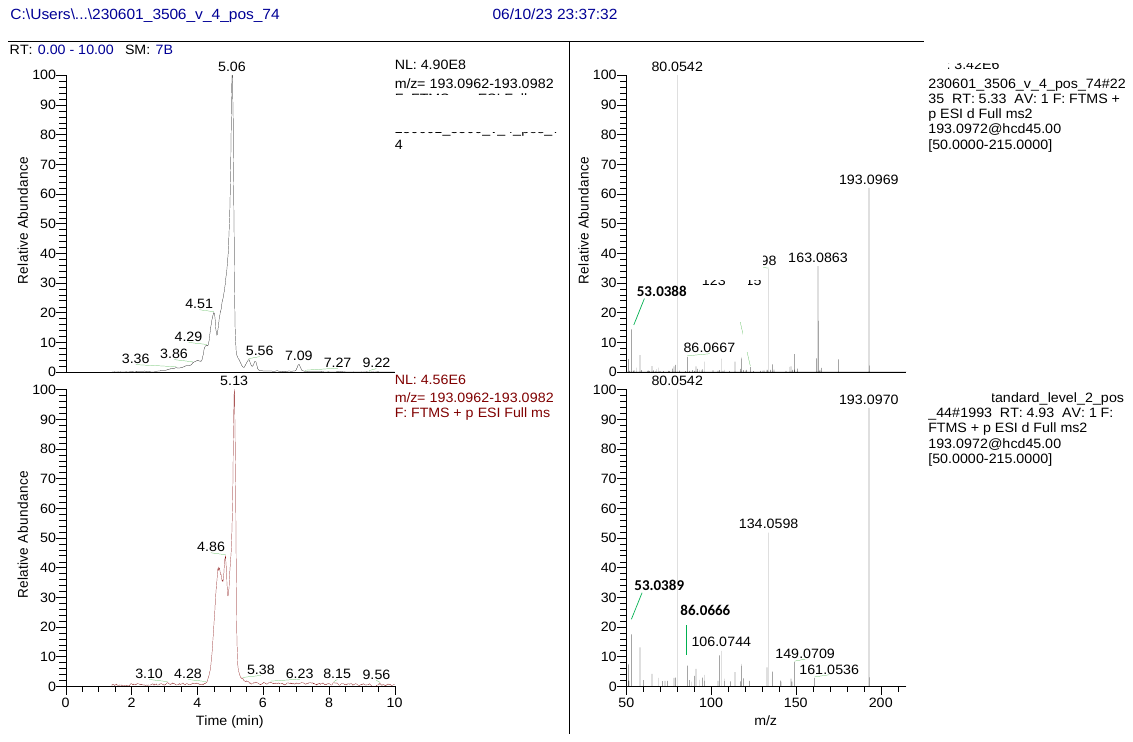

**Figure S13.** Extracted ion chromatogram (left) and data-dependent acquisition (DDA) spectrum (right) for 3-hydroxycotinine in individual plasma (top) and standard solution (bottom).

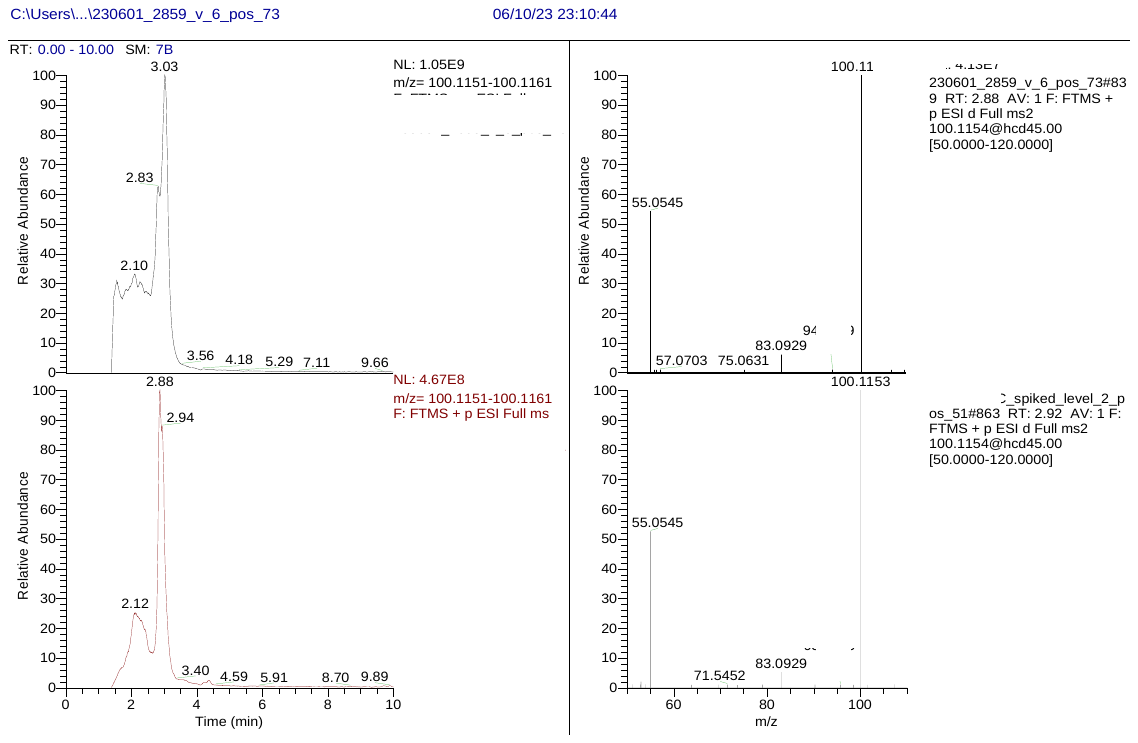

**Figure S14.** Extracted ion chromatogram (left) and data-dependent acquisition (DDA) spectrum (right) for cyclohexylamine in individual plasma (top) and standard solution (bottom).

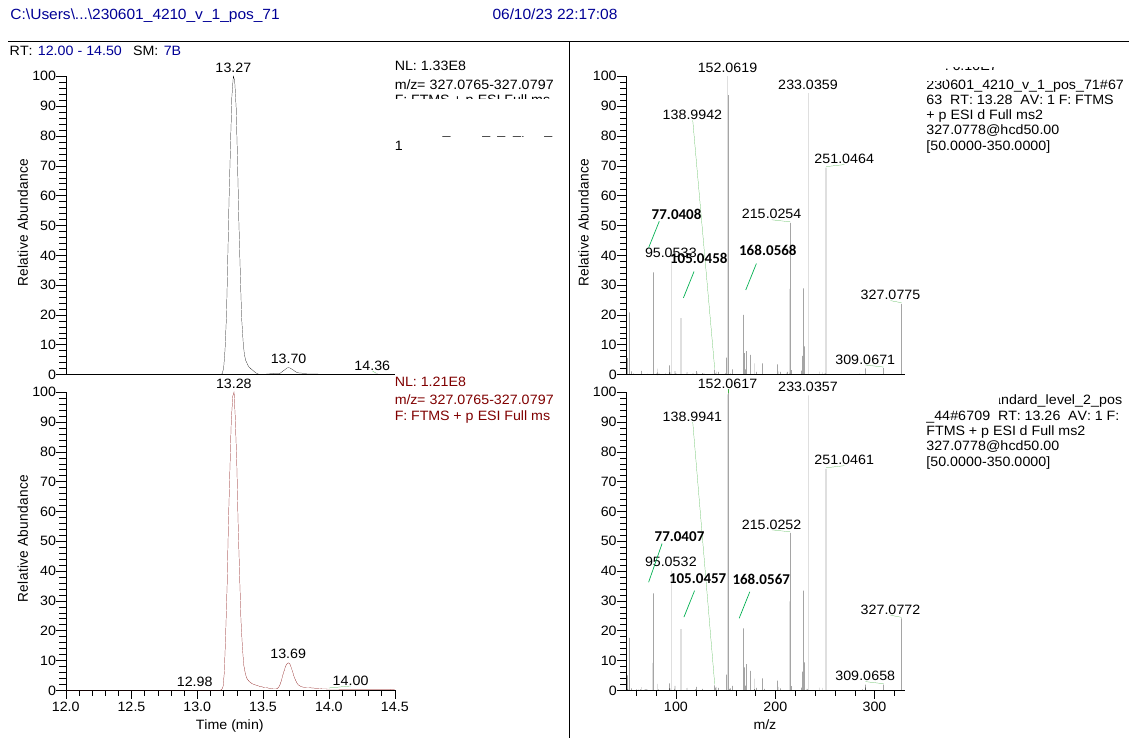

**Figure S15.** Extracted ion chromatogram (left) and data-dependent acquisition (DDA) spectrum (right) for triphenyl phosphate in individual plasma (top) and standard solution (bottom).

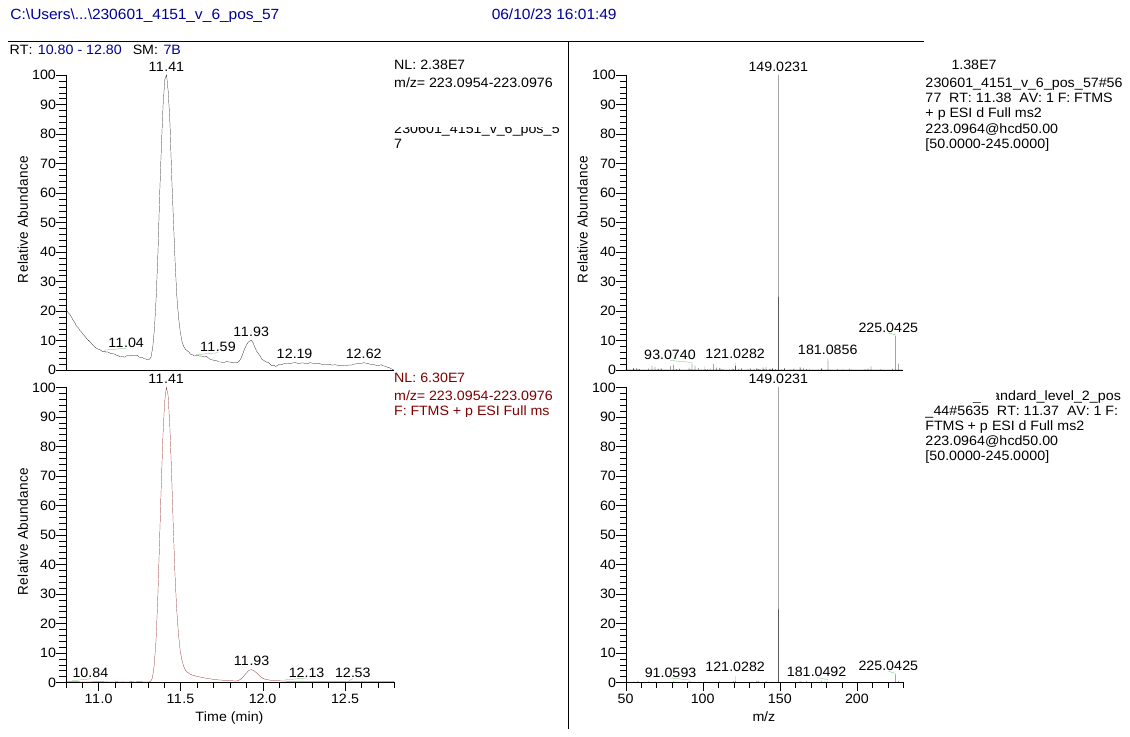

**Figure S16.** Extracted ion chromatogram (left) and data-dependent acquisition (DDA) spectrum (right) for diethyl phthalate in individual plasma (top) and standard solution (bottom).

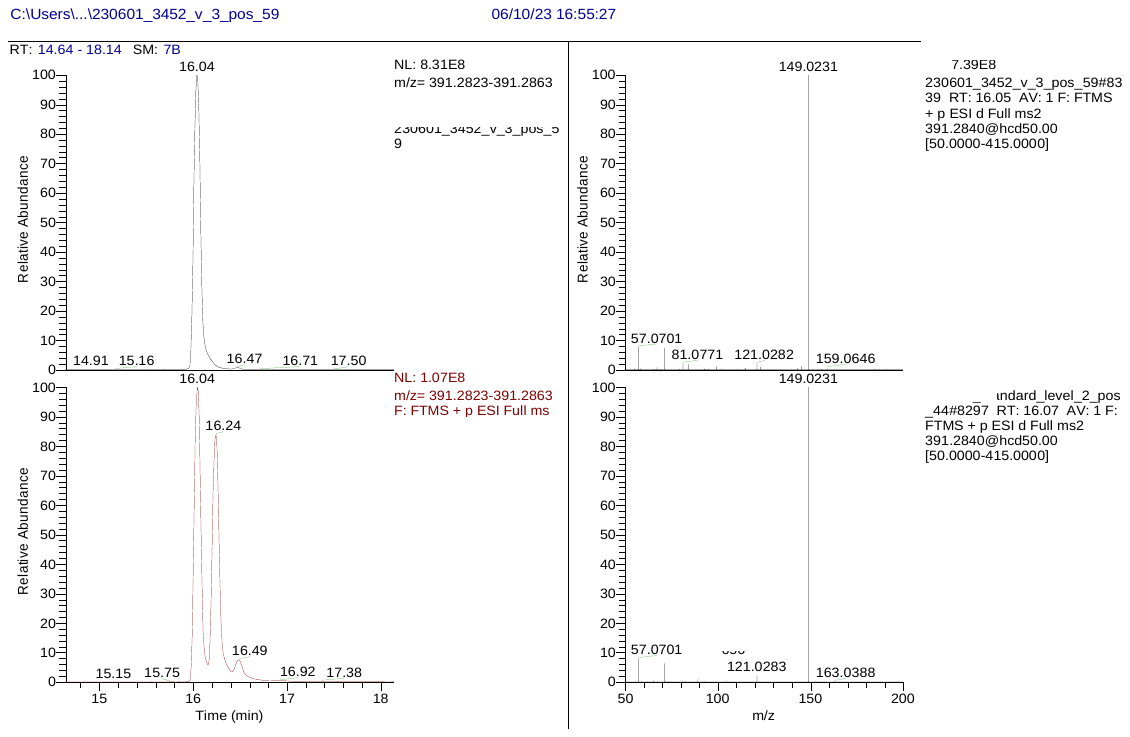

**Figure S17.** Extracted ion chromatogram (left) and data-dependent acquisition (DDA) spectrum (right) for Bis(2-ethylhexyl) phthalate (DEHP) in individual plasma (top) and standard solution (bottom). In the standard solution di-n-octyl phthalate (also spiked) elutes at 16.24 min. A background peak elutes at 16.49 in the standard solution and 16.47 in individual plasma.

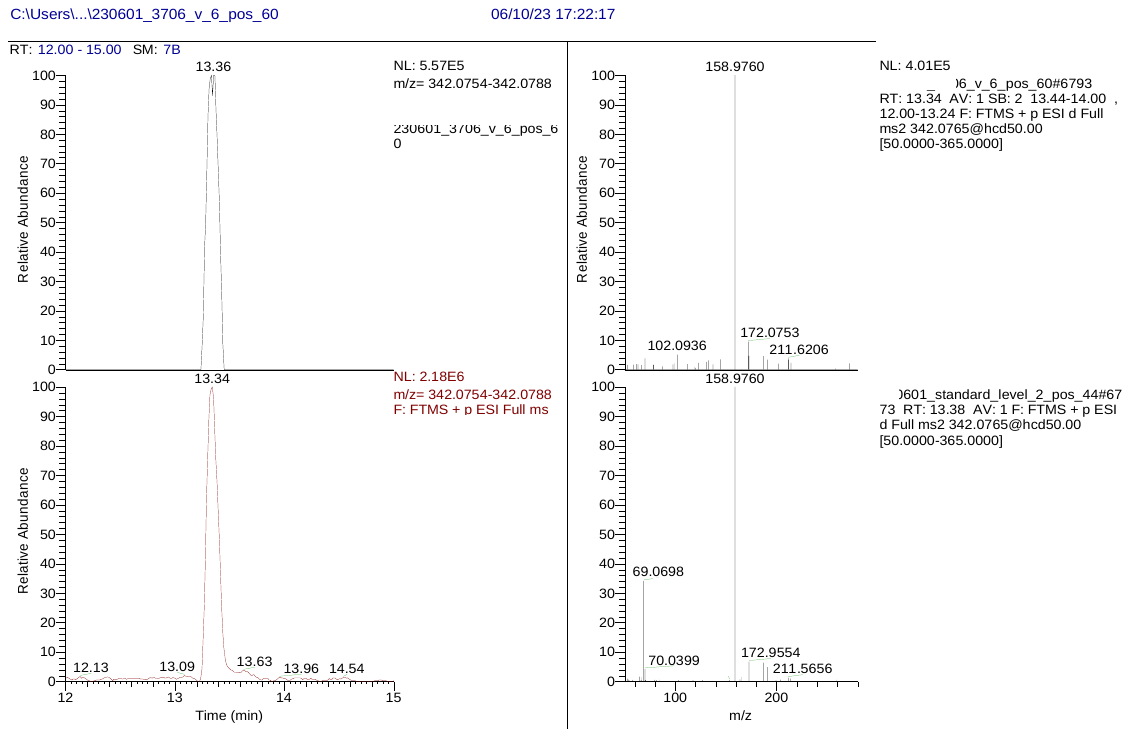

**Figure S18.** Extracted ion chromatogram (left) and data-dependent acquisition (DDA) spectrum (right) for propiconazole in individual plasma (top) and standard solution (bottom).

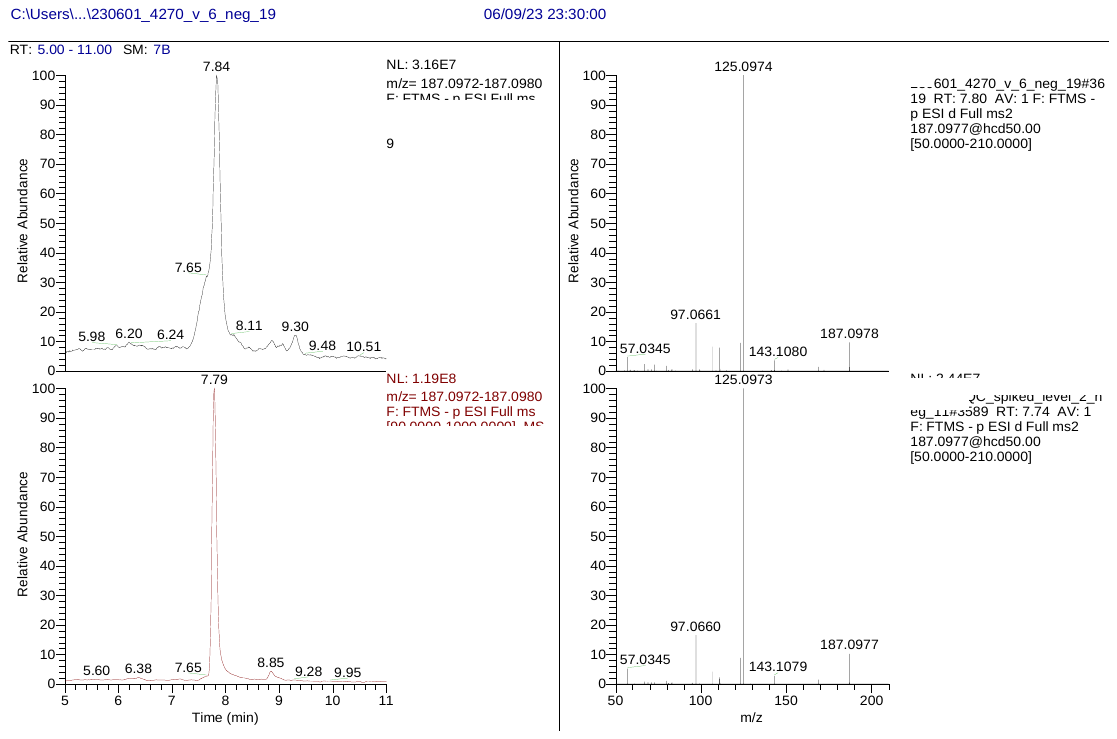

**Figure S19.** Extracted ion chromatogram (left) and data-dependent acquisition (DDA) spectrum (right) for azelaic acid in individual plasma (top) and standard solution.

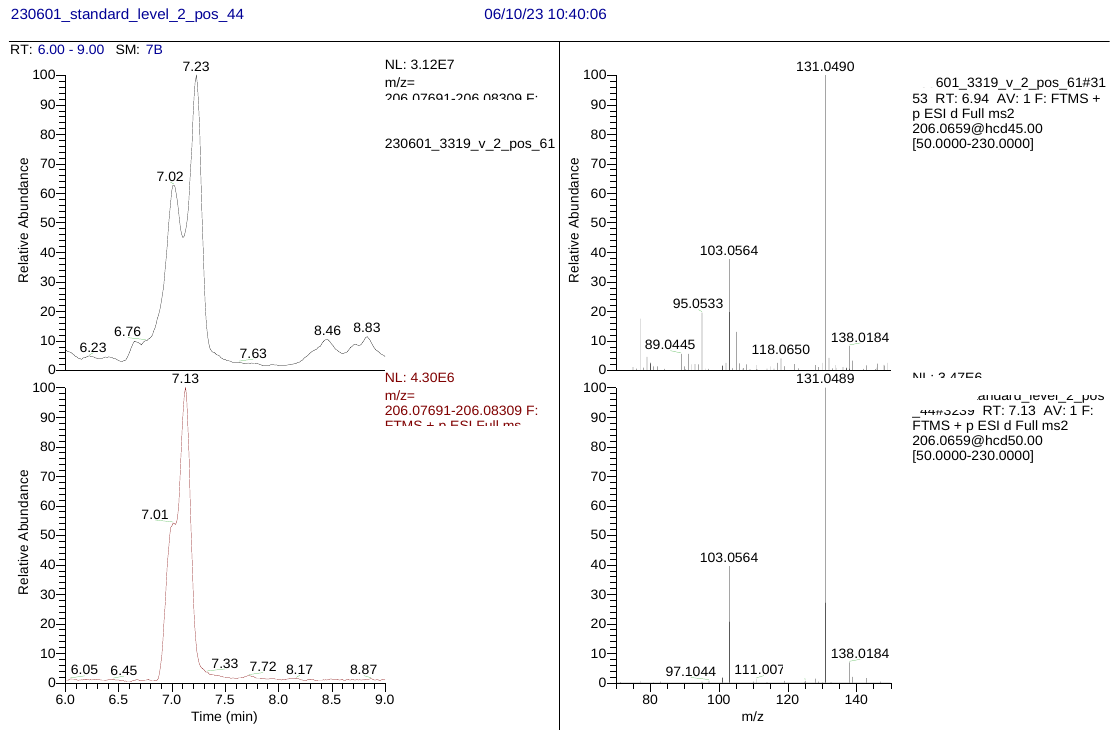

**Figure S20.** Extracted ion chromatogram (left) and data-dependent acquisition (DDA) spectrum (right) for N-cinnamoylglycine in individual plasma (top) and standard solution.

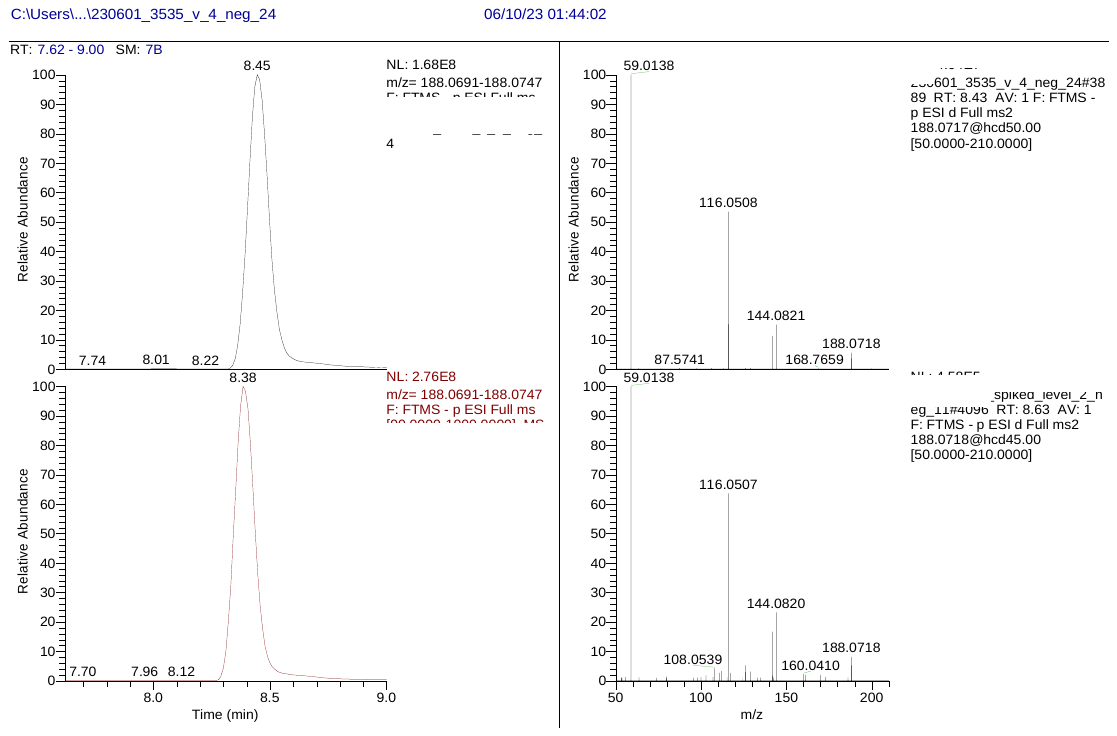

**Figure S21.** Extracted ion chromatogram (left) and data-dependent acquisition (DDA) spectrum (right) for indolepropionic acid in individual plasma (top) and standard solution.

**Figure S22.** Hierarchical cluster analysis heatmap showing exposome profiles for 46 individuals at each of 6 visits, including 519 annotated substances (Level 1 and Level 2). The entire heatmap overview is shown where color coding of features is according to class, subclass, confidence Level of identification, ionization and detection frequency (DF). Color coding of individuals is according to visit, sex, age and BMI. The order of individuals and molecular features is according to hierarchical cluster analysis conducted for the averaged responses across visits for each individual, as shown in Figure 4. A group of clustered PFAS substances is highlighted on the heatmap.

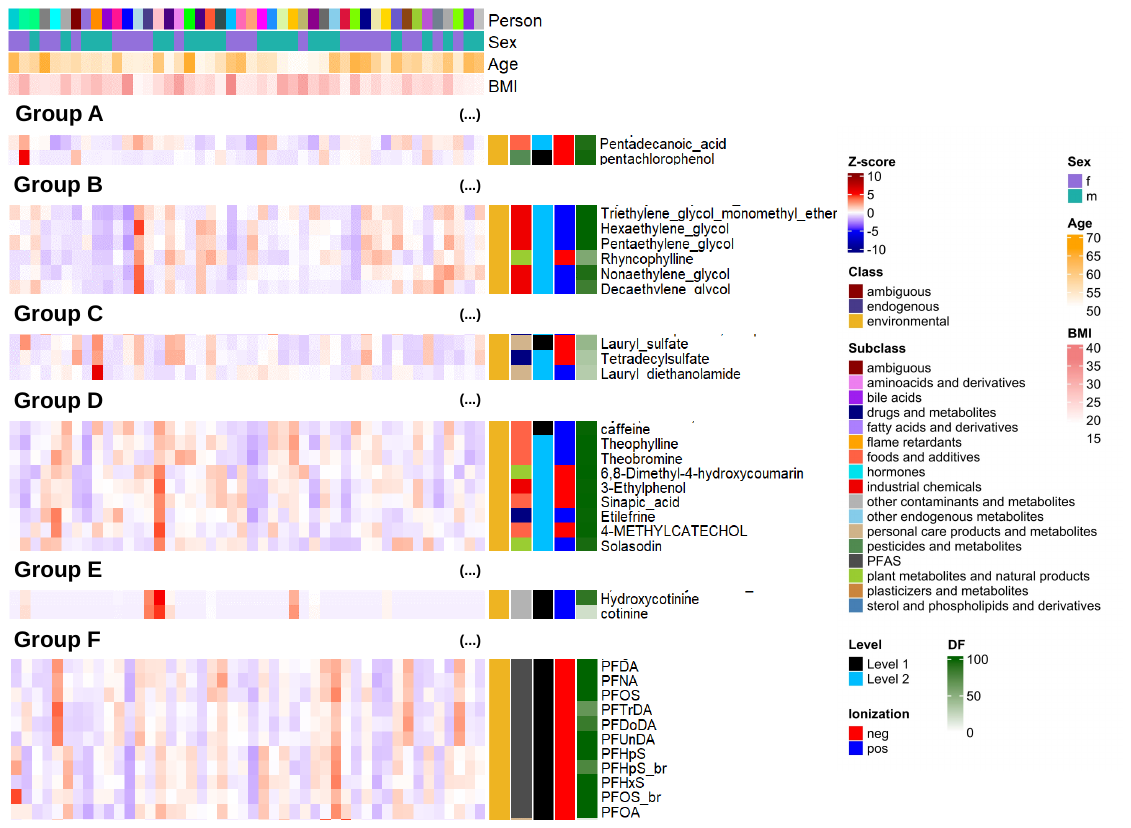

**Figure S23.** Zoomed-in sections of the hierarchical cluster analysis heatmap, showing the exposome profiles of 46 individuals, each averaged across the 6 clinical visits (full version in Figure 4). Groups of common co-exposures (detected in many participants) are shown.

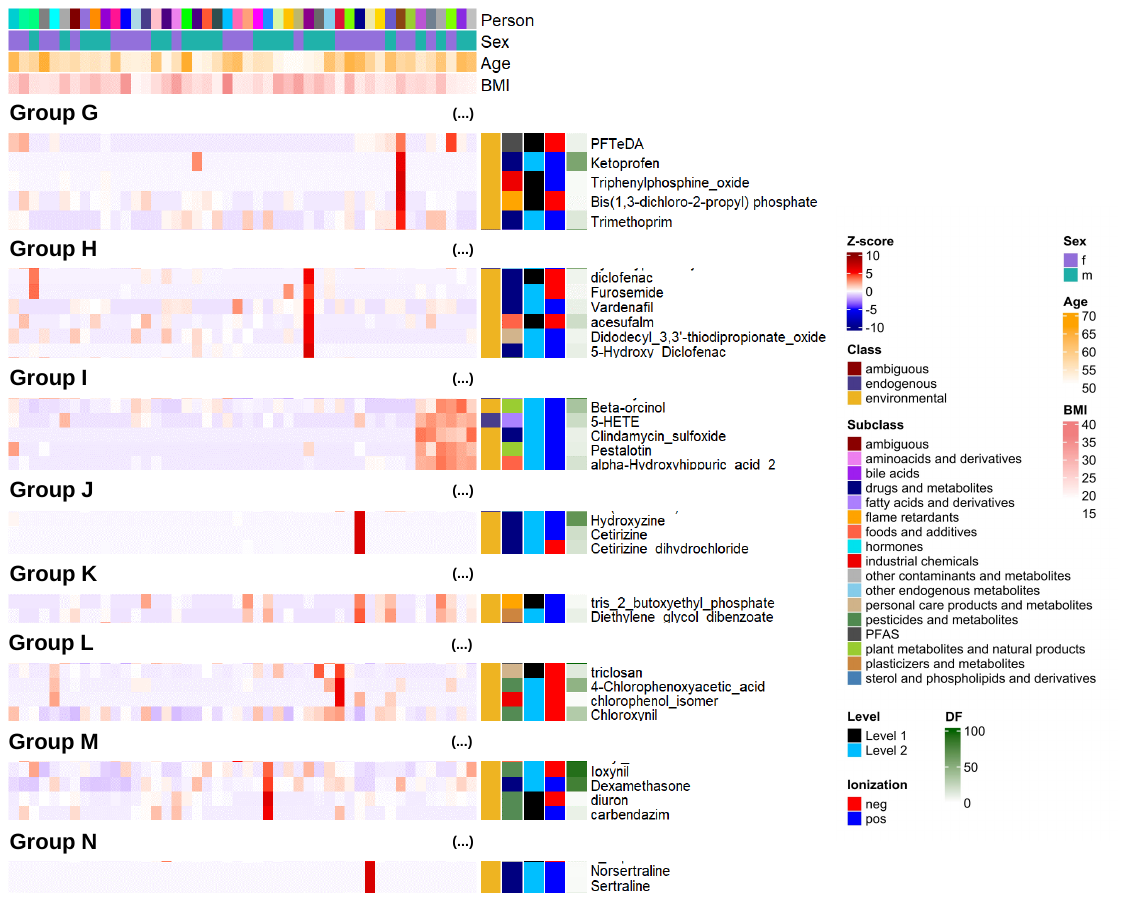

**Figure S24.** Zoomed-in sections of the hierarchical cluster analysis heatmap, showing the exposome profiles of 46 individuals, each averaged across the 6 clinical visits (full version in Figure 4). Groups of rare co-exposures (detected only in individuals or a small population fraction) are shown.

**Figure S25.** Correlations observed for PFAS targeted analytes. For linear regression the concentrations at all 6 visits of each individual (n= 276) were used and correlation was significant in all cases (p-value < 0.001). The Pearson correlation coefficient (*r*) is shown for each regression along with 95% confidence intervals and analyte distributions in histograms.

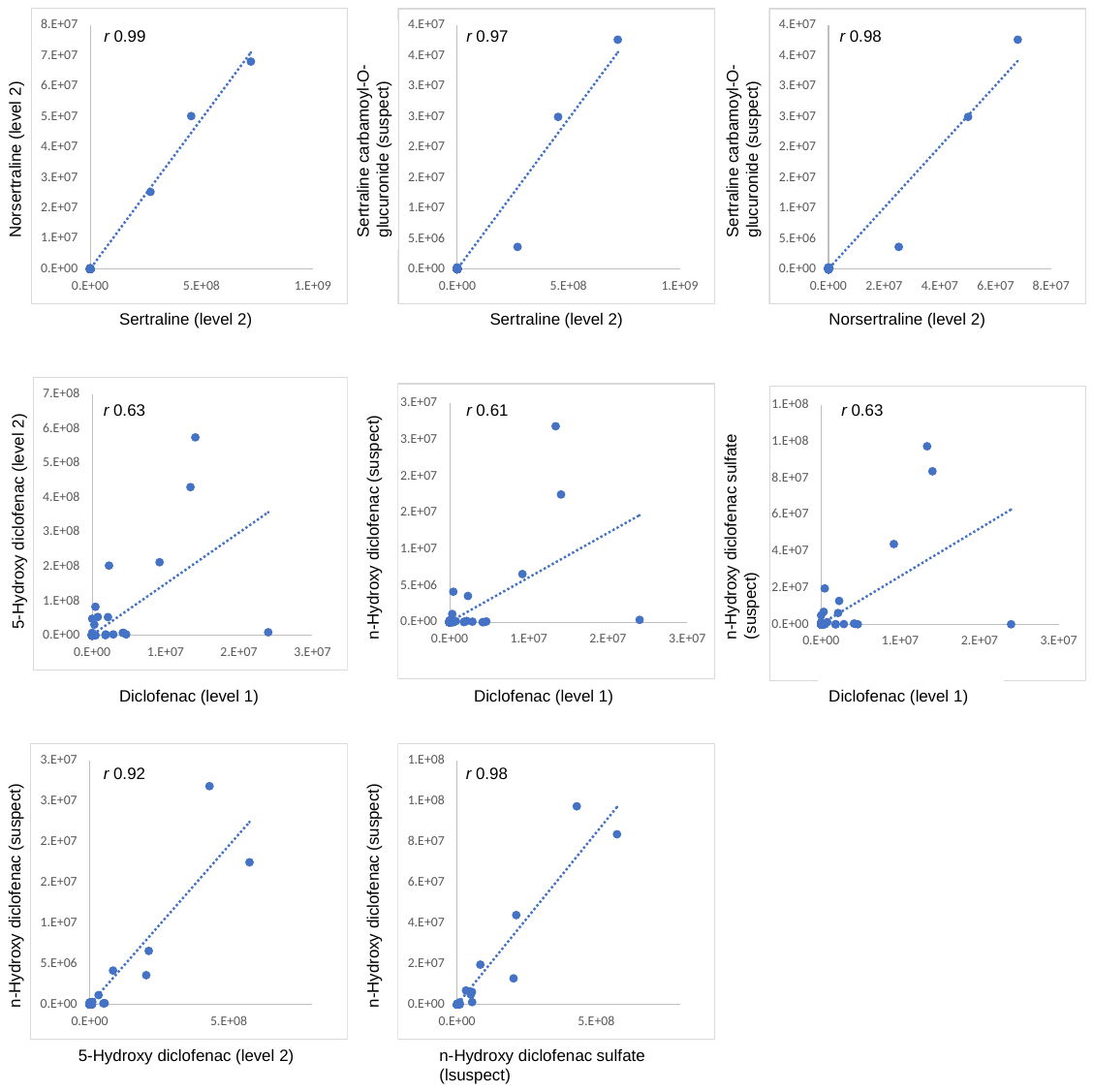

**Figure S26.** Correlations observed for Level 1 and Level 2 drugs and drug metabolites with suspect drug metabolites (not associated with a spectral library match). For linear regression the normalized areas at all 6 visits of each individual (n= 276) were used and correlation was significant in all cases (p-value < 0.001). The Pearson correlation coefficient (*r*) is shown for each regression.

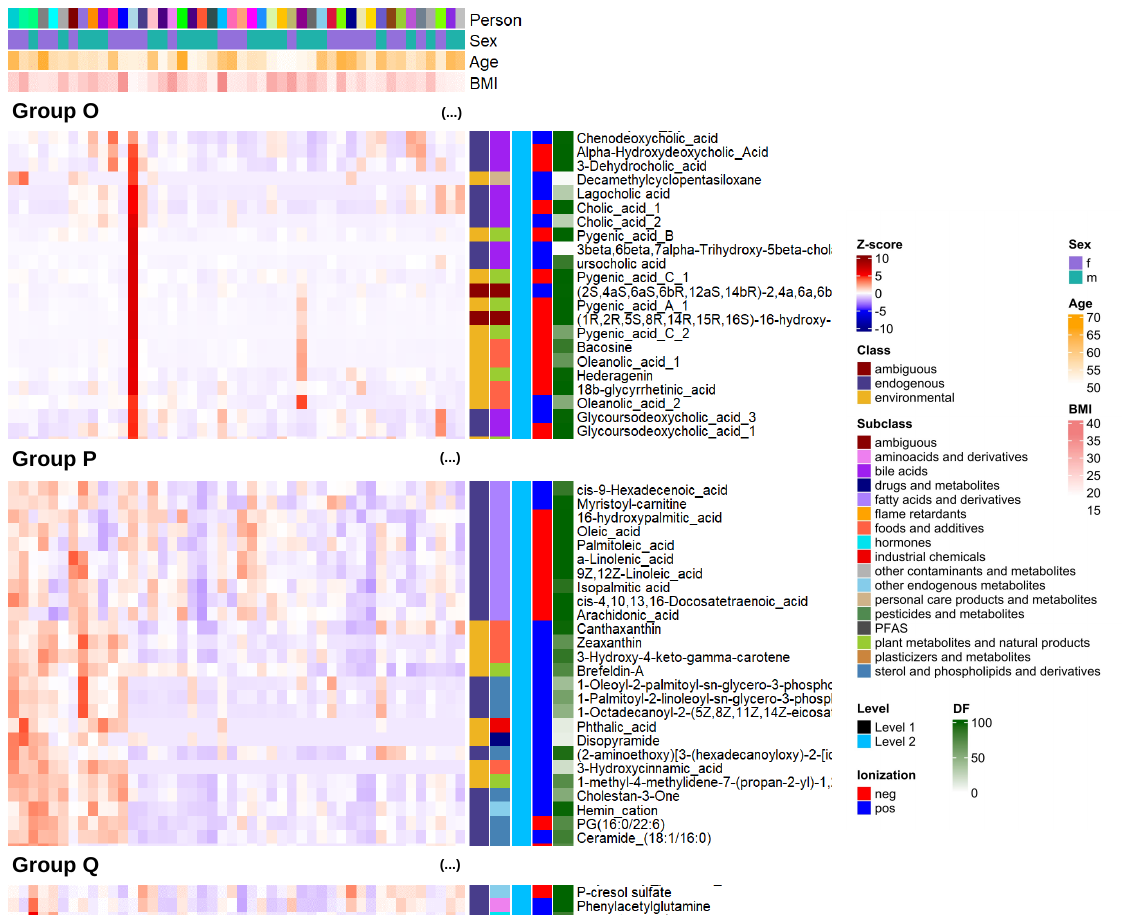

**Figure S27.** Zoomed-in sections of Hierarchical cluster analysis heatmap, showing the exposome profiles of 46 individuals, each averaged across the 6 clinical visits (full version in Figure 4). The zoomed-in sections show groups involving endogenous metabolites.

**Figure S28.** Correlations observed for Level 2 analytes classified as endogenous metabolites. Associations are shown for (A) bile acids (B) fatty acids and (C) microbial metabolites. For linear regression the normalized areas at all 6 visits of each individual (n= 276) were used and correlation was significant in all cases (p-value < 0.001). The Pearson correlation coefficient (*r*) is shown for each regression along with 95% confidence intervals and analyte distributions in histograms.

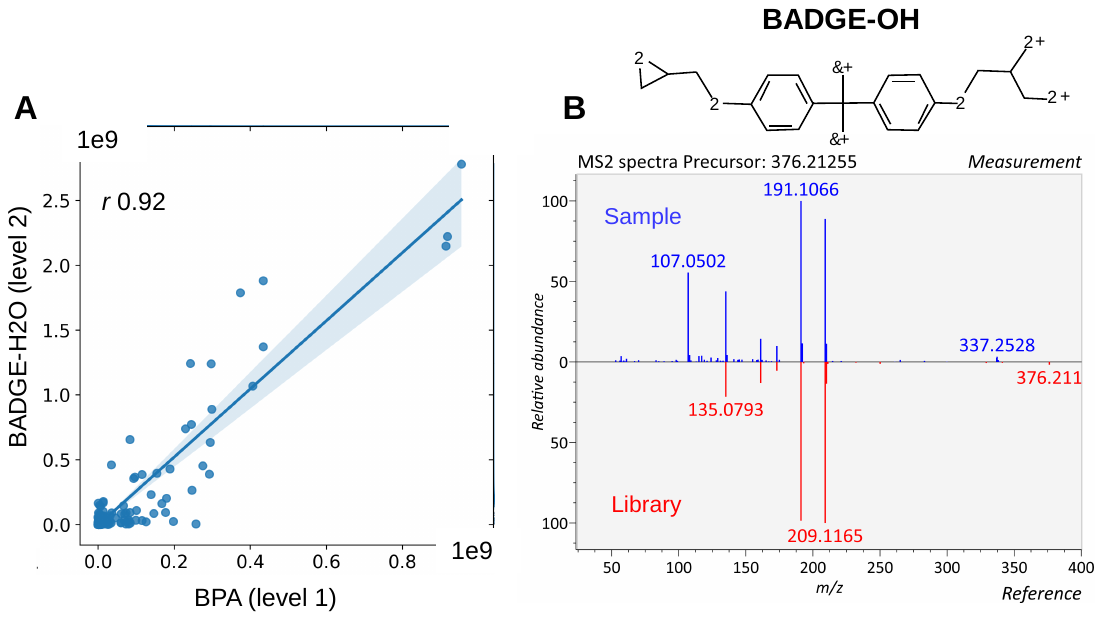

**Figure S29.** Correlation observed for BADGE-H2O annotated at Level 2 and the targeted analyte BPA. Panel (A) shows the linear regression with 95% confidence interval between BPA and BADGE-H2O, using the normalized areas at all 6 visits of each individual (n= 276). The correlation was significant (p-value < 0.001) and the Pearson correlation coefficient (*r*) is shown on panel (A). Panel (B) shows the structure for BADGE-OH and the corresponding spectral library match.

Table S1. (separate file; XSLX)

Information on labelled standard compounds used in this study.

Table S2. (separate file; XSLX)

Information on targeted analytes and corresponding native standards.

Table S3. (separate file; XSLX)

MSDIAL parameters for chromatographic alignment, spectral deconvolution and peak integration.

Table S4. (separate file; XSLX)

Targeted analytes detected and quantified in authentic human plasma samples from the S3WP cohort.

Table S5. (separate file; XSLX)

Level 1 and Level 2 annotated substances in authentic human plasma samples from the S3WP cohort.

Table S6. Discovered analytes confirmed at Level 1 with authentic standards.

| **Annotation** | **CAS-number** | **Supplier for authentic standard** | **Standard purity** |
| --- | --- | --- | --- |
| 1,3-Diphenylguanidine | 102-06-7 | AK Scientific | 98.6% |
| 1-Naphthalenesulfonic acid* | 85-47-2 | Chemtronica | 99.6% |
| 2-Naphthalenesulfonic Acid† | 76530-12-6 | Tokyo Chemical Industry (TCI) | 99.9% |
| 4-Methyl-1H-benzotriazole‡ | 29878-31-7 | Merck | 93.8% |
| 5-Methyl-1H-benzotriazole§ | 136-85-6 | Merck | 100.0% |
| 2,6-Di-tert-butyl-4-nitrophenol | 728-40-5 | Merck | NA |
| 3′-Hydroxycotinine | 34834-67-8 | Merck | 98.2% |
| 4-Chlorophenol | 106-48-9 | Merck | 99.8% |
| 4-tert-Butylpyrocatechol | 98-29-3 | Merck | 99.0% |
| Azelaic acid | 123-99-9 | Merck | 99.7% |
| Chlorothalonil-4-hydroxy | 28343-61-5 | Merck | 98.0% |
| Cyclohexylamine | 108-91-8 | VWR | 99.8% |
| Sodium lauryl sulfate | 151-21-3 | Merck | 98.7% |
| Propiconazole | 60207-90-1 | Honeywell | 98.3% |
| Triphenyl phosphate | 115-86-6 | Merck | 99.8% |
| Triphenylphosphine oxide | 791-28-6 | Merck | 99.8% |
| Indolepropionic acid | 830-96-6 | Merck | 99.9% |
| N-cinnamoylglycine | 16534-24-0 | Merck | 100.0% |
| Diethyl phthalate | 84-66-2 | LGC | 98.7% |
| Di-2-ethylhexyl phthalate (DEHP) | 117-81-7 | LGC | 99.7% |

* cointegrated together with 2-Naphthalenesulfonic Acid as "1- / 2-naphthalenesulfonate (mix)"

†cointegrated together with 1-Naphthalenesulfonic Acid as "1- / 2-naphthalenesulfonate (mix)"

‡coelution with 5-methyl-1H-benzotriazole

§coelution with 4-methyl-1H-benzotriazole

NA: not available

Table S7. (separate file; XSLX)

Intraclass correlation coefficient (ICC) of Level 1 and Level 2 annotated substances.

Table S8. Mixed-effects model parameters for the associations between female testosterone (dependent variable, log ng/L) and PFAS targeted analytes (independent variable, log ng/L). For each PFAS, the parameters (Y-intercepts, slope (β)) of the unadjusted models are shown in the first row along with associated p-values (raw and Bonferroni adjusted). In the second row for each PFAS, the corresponding adjusted model is shown that controls for both baseline age and body mass index (BMI) as fixed effects.

| **PFAS / Model** | ***Y-int*** | ***β*** | ***p (raw)* sig/non-sig** | ***p (Bonferroni)* sig/non-sig** |
| --- | --- | --- | --- | --- |
| **PFHxS** | -0.75524 | 0.39445 | 0.00845 | 0.10135 |
| -Adjusted for BMI+Age | -0.34530 | 0.37940 | 0.01780 | 0.21360 |
| **br-PFHxS** | -0.79545 | -0.00176 | 0.98091 | 1.00000 |
| -Adjusted for BMI+Age | -0.52074 | -0.02085 | 0.78497 | 1.00000 |
| **PFHpS** | -0.65583 | 0.15354 | 0.22804 | 1.00000 |
| -Adjusted for BMI+Age | -0.06756 | 0.14385 | 0.27416 | 1.00000 |
| **br-PFHpS** | -0.76912 | 0.01777 | 0.72125 | 1.00000 |
| -Adjusted for BMI+Age | -0.20121 | 0.01283 | 0.79917 | 1.00000 |
| **PFOS** | -1.07134 | 0.40605 | 0.00059 | 0.00714 |
| -Adjusted for BMI+Age | -0.46000 | 0.37660 | 0.00163 | 0.01957 |
| **br-PFOS** | -0.86803 | 0.27146 | 0.23480 | 1.00000 |
| -Adjusted for BMI+Age | 0.00033 | 0.21268 | 0.14149 | 1.00000 |
| **PFHpA** | -0.81174 | -0.01541 | 0.65005 | 1.00000 |
| -Adjusted for BMI+Age | -0.49248 | -0.02026 | 0.55531 | 1.00000 |
| **PFOA** | -0.87362 | 0.33615 | 0.04522 | 0.54263 |
| -Adjusted for BMI+Age | -0.43620 | 0.29282 | 0.09001 | 1.00000 |
| **PFNA** | -0.72828 | 0.54124 | 0.00006 | 0.00069 |
| -Adjusted for BMI+Age | -0.66928 | 0.46864 | 0.00754 | 0.09042 |
| **PFDA** | -0.59725 | 0.48636 | 0.00002 | 0.00019 |
| -Adjusted for BMI+Age | -0.62981 | 0.47440 | 0.00003 | 0.00030 |
| **PFUnDA** | -0.65593 | 0.34086 | 0.01037 | 0.12441 |
| -Adjusted for BMI+Age | -0.41643 | 0.33229 | 0.01824 | 0.21891 |
| **PFTrDA** | -0.69053 | 0.07563 | 0.03753 | 0.45040 |
| -Adjusted for BMI+Age | -0.45202 | 0.10241 | 0.01014 | 0.12165 |
